# Structural insights into the recognition of mono- and di-acetylated histones by the ATAD2B bromodomain

**DOI:** 10.1101/263624

**Authors:** Jonathan T. Lloyd, Kyle McLaughlin, Mulu Y. Lubula, Jamie C. Gay, Andrea Dest, Cong Gao, Margaret Phillips, Marco Tonelli, Gabriel Cornilescu, Matthew R. Marunde, Chiara M. Evans, Samuel P. Boyson, Samuel Carlson, Michael-Christopher Keogh, John L. Markley, Seth Frietze, Karen C. Glass

**Affiliations:** Department of Pharmaceutical Sciences, Albany College of Pharmacy and Health Sciences, Colchester, VT, 05446, USA; Department of Biomedical and Health Sciences, University of Vermont, Burlington, VT 05405, USA; National Magnetic Resonance Facility at Madison and Department of Biochemistry, University of Wisconsin-Madison, Madison, WI, 53706, USA; EpiCypher Inc, Durham, NC, USA

**Keywords:** Acetyllysine, bromodomain, chromatin reader domain, epigenetics, histone, post-translational modification

## Abstract

Bromodomains are chromatin reader modules that recognize acetylated lysine. Different bromodomains exhibit a preference for specific patterns of lysine acetylation marks on core and variant histone proteins, however, the functional relationships that exist between histone acetyllysine ligands and bromodomain recognition remain poorly understood. In this study, we examined the ligand specificity of the ATAD2B bromodomain and compared it to its closely related paralog in ATAD2. We show that the ATAD2B bromodomain selects for mono- and di-acetylated histones, and structural analysis identified key residues in the acetyllysine binding pocket that dictate ligand binding specificity. The X-ray crystal structure of the ATAD2B bromodomain in complex with an ATAD2 bromodomain inhibitor was solved at 2.4 Å resolution. This structure demonstrated that critical contacts required for bromodomain inhibitor coordination are conserved between the ATAD2/B bromodomains, and many of these residues play a dual role in acetyllysine recognition. We further characterized a variant of the ATAD2B bromodomain that through alternative splicing loses critical amino acids required for histone ligand and inhibitor coordination. Altogether our results outline the structural and functional features of the ATAD2B bromodomain and identify a novel mechanism important for regulating the interaction of the ATAD2B protein with chromatin.

**HIGHLIGHTS:**

- The ATAD2B bromodomain recognizes mono- and di-acetylated histone ligands.
- Chemical shift perturbations outline the ATAD2B bromodomain acetyllysine binding pocket.
- An ATAD2B bromodomain-inhibitor complex reveals important binding contacts.
- An alternate splice variant in the ATAD2B bromodomain abolishes histone and inhibitor binding.

## INTRODUCTION

Post-translational modifications (PTMs) on histone proteins play a central role in the regulation of chromatin structure. Lysine acetylation (Kac) is a major PTM widely studied in the establishment of cell-type specific enhancer patterns. Although largely associated with the opening of chromatin structure and activation of gene transcription^1–3^, recent studies have found that specific Kac histone marks regulate chromatin compaction and DNA damage responses^4,5^. Altered histone acetylation patterns have been linked to disease progression, which may arise due to the deregulation of enzymes responsible for adding or removing Kac marks, or through the protein interaction modules that recognize and interpret Kac.

Bromodomains are evolutionarily conserved protein−protein interaction modules that serve as Kac epigenetic reader domains. Bromodomain-containing proteins interact with transcription factor and chromatin-modifying complexes and have been reported to participate in epigenetic memory^6^. In humans, there are 61 known bromodomains found in 46 different proteins^7^. These bromodomains are divided into eight subfamilies (I-VIII) based on sequence and structural features^7^. Most families contain a canonical bromodomain that exhibits binding preference for different histone Kac ligands on core and variant histone proteins. However, atypical bromodomains, such as the TRIM28/KAP1 bromodomain in family VI, lack essential acetyllysine recognition residues and display little to no affinity towards Kac-containing histone peptides^7^. These non-canonical bromodomains may serve to recognize alternative histone PTMs^7,8^, or they may interact with acetylation marks found on non-histone proteins^9^. Interestingly, some bromodomains, including the family IV bromodomains in BRD7/9, can accommodate larger acyl groups in their binding pockets and have been shown to recognize butyrylated and crotonylated lysine^10^.

The Kac binding activity of the family IV bromodomain-containing proteins, which include the bromodomain-containing proteins ATAD2, ATAD2B, BRPF1, BRD1, BRPF3, BRD7, and BRD9, has been at least partially characterized using high throughput histone peptide arrays and biochemical and biophysical approaches (see Lloyd et al., 2017 for a recent review)^11^. The family IV bromodomains recognize mono- and di-acetylated lysine residues on histones H2A, H3, and H4. Notably, there is currently no ligand information available for the ATPase family, AAA^+^ domain-containing protein 2B (ATAD2B, also known as KIAA1240). ATAD2B is a nuclear protein that contains two conserved domains: an AAA^+^ ATPase domain and a bromodomain^12^. The AAA^+^ or ‘ATPases associated with diverse cellular activities’ domain is found in a large superfamily of proteins that typically form a hexameric complex, able to recognize ATP in order to drive molecular remodeling reactions^13^. The C-terminal bromodomain of ATAD2B is highly homologous to the ATAD2 bromodomain, with 74.7% sequence identity and 94.4% amino-acid similarity^1^. The ATAD2 bromodomain binds to acetylated histones at H4K5ac and H4K12ac, and X-ray crystal structures have elucidated essential features of the binding pocket important for ligand coordination^14,15^. Recently, the ATAD2 bromodomain was shown to recognize di-acetylated histones, and it associates with H4K5acK12ac modifications found on newly synthesized histones following DNA replication^16^. However, the acetylated histone ligands of the ATAD2B bromodomain have yet to be determined, and the molecular mechanism(s) underlying acetyllysine recognition is currently unclear.

We hypothesized that the ATAD2B bromodomain is likely involved in the recognition of di-acetyllysine modifications on the histone tail, similar to its ATAD2 paralog^16^. To outline the function of the ATAD2B bromodomain, we used a newly developed peptide array technology to screen against a large number of acetylated histone tail peptides, and characterized their binding affinity and specificity determinants with a combination of isothermal titration calorimetry (ITC) and nuclear magnetic resonance (NMR) experiments. We determined that the ATAD2B bromodomain recognizes acetylated lysine residues on the N-terminal tails of histones H4 and H2A. ITC was used to characterize the binding affinities of the ATAD2B bromodomain with histone peptide ligands, and we found that histone H4K5ac (1-15) ligand was the strongest binder. Since the molecular details of acetyllysine recognition have not been determined, we used NMR chemical shift perturbation data to map out the Kac binding pocket on the apo ATAD2B bromodomain structure (PDB ID: 3LXJ). This allowed us to identify the amino acid residues lying within and outside of the canonical Kac binding site that are affected by the addition of the acetyllysine ligands. Site-directed mutagenesis coupled to ITC experiments confirmed the role of residues within the bromodomain-binding pocket in ligand coordination. X-ray crystallography was used to solve the structure of the ATAD2B bromodomain in complex with compound 38, an ATAD2 bromodomain inhibitor developed by GlaxoSmithKline^17^. From this structure, we identified the molecular contacts important for inhibitor coordination that are conserved between the ATAD2/B bromodomains, many of which are also involved in Kac recognition. Our research supports a model whereby inherent flexibility in the ATAD2/B bromodomain ZA and BC loop regions is important for recruiting Kac ligands into the bromodomain binding pocket. Notably, we describe a novel ATAD2B bromodomain isoform that lacks the ZA loop in the bromodomain Kac binding pocket that completely abrogates binding to both inhibitor and histone ligands. Interestingly, this splice variant is not observed in the ATAD2 bromodomain. Our results establish key differences between the ATAD2/B bromodomains, suggesting that these proteins carry out divergent functions within the cell. These data advance our understanding of how the ATAD2B bromodomain recognizes and selects for specific Kac modifications and provides new information that will inform the design of small molecule compounds to specifically select for the ATAD2/B bromodomains.

## RESULTS

### The ATAD2B bromodomain recognizes mono- and di-acetylated histones

Bromodomains are known as Kac reader domains^18^, however, the histone ligands of the ATAD2B bromodomain have yet to be determined. To identify the Kac modifications recognized by the ATAD2B bromodomain, we used a combination of peptide array, NMR, and ITC methods. We first used the dCypher platform to screen the recombinant ATAD2 and ATAD2B bromodomains against 288 histone peptides (containing single or combinatorial PTMs) after determining optimal protein concentrations (**Supplementary Fig. 1A-B)**. The PTMs tested included lysine methylation (me1-2-3), arginine methylation (me1-2a-2s), lysine acylation (acetyl, butyryl, crotonyl, propionyl), and phosphorylation (on S/T/Y). The ATAD2B bromodomain bound to 39 PTM peptide ligands from histones H4, H2A, and the histone variant H2A.X (**Fig. 1A-B**); predominantly recognizing acetyllysine modifications, but permissive for adjacent PTMs (e.g. methylation, phosphorylation) or alternate short-chain acylations (propionylation and butyrylation). In comparison, the ATAD2 bromodomain appears to recognize a more limited subset of histone ligands, as significant binding interactions were only observed with 11 peptide ligands on histone H4. To identify the preferred ligands of the ATAD2B bromodomain we further characterized their binding affinity using ITC (**Table 1, Fig. 1C, and Supplementary Fig. 2**). We discovered that the ATAD2B bromodomain selects for both mono- and di-acetylated histone ligands, preferentially binding to the histone H4K5ac (1-15) peptide with the strongest affinity (K_D_= 5.2 ± 1.0 μM), followed by recognition of several di-acetylated ligands including H4K5acK12ac (1-15) and H4K5acK8ac (1-10). The weak interaction of the ATAD2B bromodomain with H4K8ac (1-10) (K_D_= 1164.2 ± 28.5 μM) (**Fig. 1C**), was confirmed by NMR chemical shift perturbations observed in the bromodomain binding pocket upon addition of this ligand (**Supplementary Fig. 3A**). We also found that the first three residues of the histone H4 tail are crucial for acetyllysine recognition by the ATAD2B bromodomain, especially when the acetylation mark occurs on the K5 or K8 residue (**Table 1**). Altogether our data indicate that the ATAD2B bromodomain preferentially selects for mono- and di-acetylated histone ligands on histones H4 and H2A, and its binding pocket spans at least 10 amino acid residues in order to recognize histone ligands H4K5acK12ac and H4K5acK16ac.

**Table 1.** Binding of the ATAD2B bromodomain to acetylated histones. Dissociation constants of the interaction of histone tail peptides with the ATAD2B bromodomain measured by ITC. Sequences of the acetylated histone peptides are also indicated.

| Histone peptide | Sequence | ATAD2B<br>ITC $K_D$ (μM) | ATAD2A<br>Reported $K_D$s |
| --- | --- | --- | --- |
| H2AK5ac (res 1-12)* | SGRG(Kac)QGGKARA | $42.5 \pm 2.9$ | NA |
| H4 unmodified (res 4-17)* | GKGGKGLGKGGAKRHRK | No Binding | NA |
| H4K5ac (res 1-15)* | SGRG(Kac)GGKGLGKGGA | $5.2 \pm 1.0$ | NA |
| H4K5ac (res 1-10)* | SGRG(Kac)GGKGL | $48.0 \pm 4.7$ | 22 μM<br>(res 1-20)<sup>14</sup> |
| H4K5ac (res 4-17)* | G(Kac)GGKGLGKGGAKR | No Binding | NA |
| H4K8ac (res 1-10)* | SGRGKGG(Kac)GL | $1164.2 \pm 28.5$ | NA, interaction shown<sup>16</sup> |
| H4K8ac (res 4-17) | GKGG(Kac)GLGKGGAKR | No Binding | NA |
| H4K12ac (res 4-17)* | GKGGKGLG(Kac)GGAKR | $50.5 \pm 5.8$ | 2.5 μM<br>(res 1-25)<sup>16</sup> |
| H4K12ac (res 1-15) | SGRGKGGKGLG(Kac)GGA | $685.3 \pm 91.7$ | NA |
| H4K16ac (res 10-20)* | LGKGGA(Kac)RHRK | No Binding | NA |
| H4K5acK8ac (res 1-10)* | SGRG(Kac)GG(Kac)GL | $28.1 \pm 1.6$ | NA |
| H4K5acK12ac (res 1-15)* | SGRG(Kac)GGKGLG(Kac)GGA | $18.7 \pm 0.9$ | NA, interaction shown<sup>16</sup> |
| H4K5acK16ac (1-24) | SGRG(Kac)GGKGLGKGGA(Kac)<br>RHRKVLRD | $34.4 \pm 6.2$ | NA |
| H4K8acK12ac (res 6-15) | GG(Kac)GLG(Kac)GGA | No Binding | NA |
| H4K8acK16ac (1-20) | SGRGKGG(Kac)GLGKGGA(Kac)<br>RHRK | $35.7 \pm 6.5$ | NA |
| H4K12acK16ac (res 10-20) | LG(Kac)GGA(Kac)RHRK | $213.9 \pm 7.0$ | NA |
| H3 unmodified (res 1-12)* | ARTKQTARKSTG | No Binding | NA |
| H3K9ac (res 1-12) | ARTKQTAR(Kac)STG | No Binding | NA |
| H3K14ac (res 8-19)* | RKSTGG(Kac)APRKQ | No Binding | No binding<br>(res 1-21)<sup>14</sup> |
| Compound 38 | --- | $160.6 \pm 12.3$<br>(nM) | 90 (nM)<sup>17</sup> |

\*Indicates peptides tested in NMR chemical shift perturbation experiments (Figure 2B and Supplemental Figure 2A)

**Figure 1.**
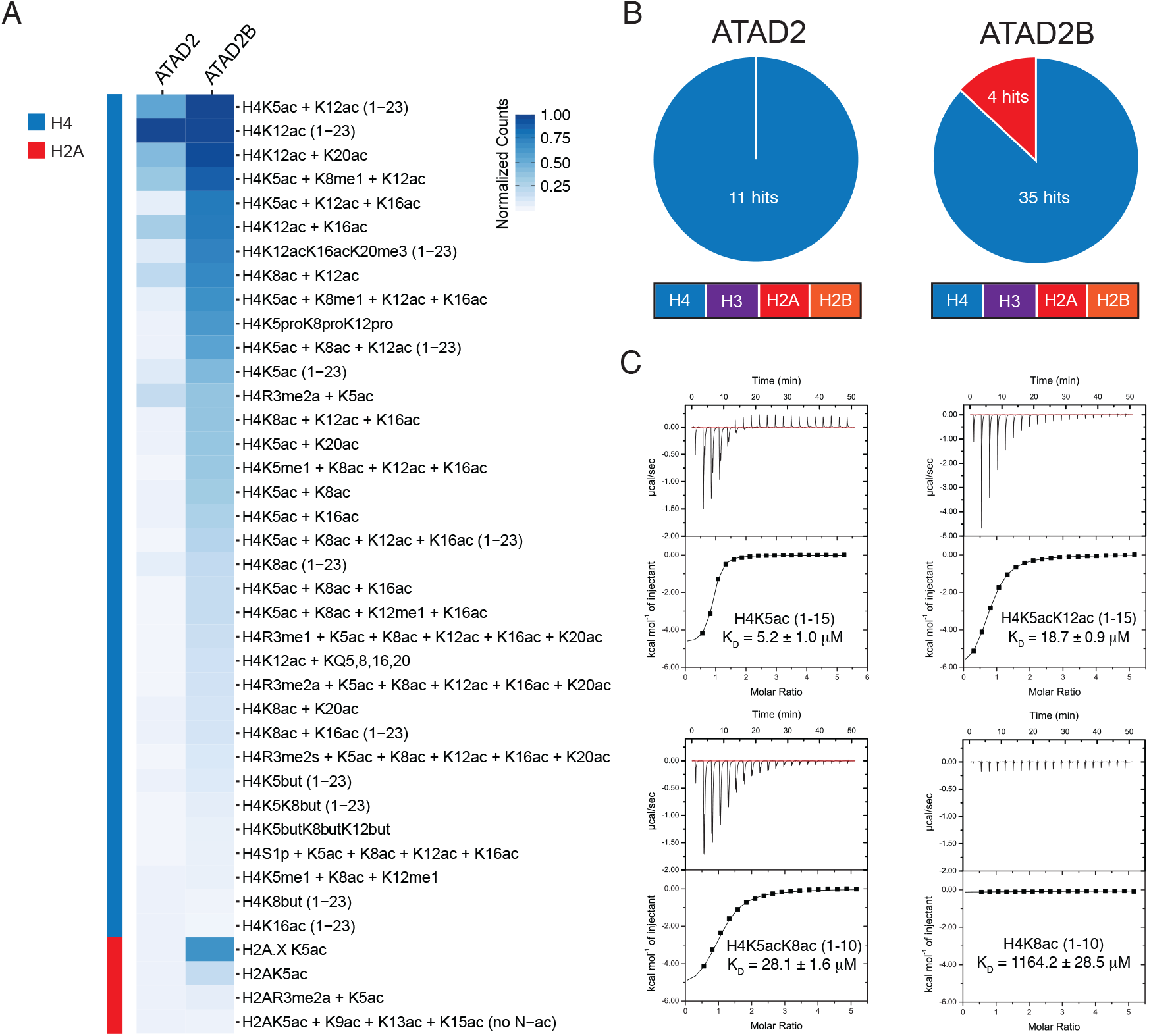
The ATAD2B bromodomain recognizes acetylated histones. (A) Heat map represents relative binding of GST-tagged ATAD2 and ATAD2B bromodomains to histone peptides using AlphaScreen technology (i.e. dCypher phase B: see Methods). Alpha counts (n=2) were normalized for each protein to the highest fluorescent intensity signal for each respective assay, and the relative binding strength is indicated by the color gradient. (B) The pie charts for ATAD2/B bromodomains indicate the number of peptides classed as positive binders (alpha counts having signals >5000, which is at least twice that of the relevant unmodified control peptide) (full dCypher peptide screen data in **Supplemental Tables 1 & 2, and Resources Table A**). (C) Exothermic ITC enthalpy plots for the binding of the ATAD2B bromodomain to mono- and di-acetylated histone ligands. The calculated binding constants are indicated.

### Mapping of the ATAD2B bromodomain binding pocket

The X-ray crystal structure of the apo ATAD2B bromodomain has been solved (PDB ID: 3LXJ), but there is no data available on the molecular mechanism of histone recognition. To obtain specific information on how the ATAD2B bromodomain recognizes histone ligands at the molecular level we first completed the NMR backbone assignment of the ^15^N,^13^C-labeled ATAD2B bromodomain (**Supplementary Fig. 3B**). To outline the Kac binding pocket within the ATAD2B bromodomain we used 2D ^1^H-^15^N HSQC NMR titration experiments involving the addition of unlabeled, mono- and di-acetylated histone ligands to the ^15^N labeled ATAD2B bromodomain protein (**Fig. 2**). Residues Lys 985, Val 987, Glu 991, Glu 997, Val 998, Thr 1007, Tyr 1037, Asn 1038, Asp 1042, Asp 1045, Arg 1051 and Leu 1055 in the ATAD2B bromodomain consistently exhibited the most significant chemical shift perturbations (CSPs) upon addition of the histone H4K5ac (1-15), H4K5acK8ac (1-10) and H4K5acK12ac (1-15) ligands. These ligands also showed the highest binding affinity as observed by ITC (**Table 1**). Of these residues, Val 987 lines the hydrophobic Kac binding pocket. and Tyr 1037 lies adjacent to Asn 1038, which is the universally conserved asparagine known to coordinate the Kac moiety^19^. Residues Glu 991 and Glu 997 reside in the ZA loop. Residues Thr 1007, Arg 1051 and Leu 1055 are present in the αC helix, while residues Asp 1042 and Asp 1045 are located in the flexible BC loop.

**Figure 2.**
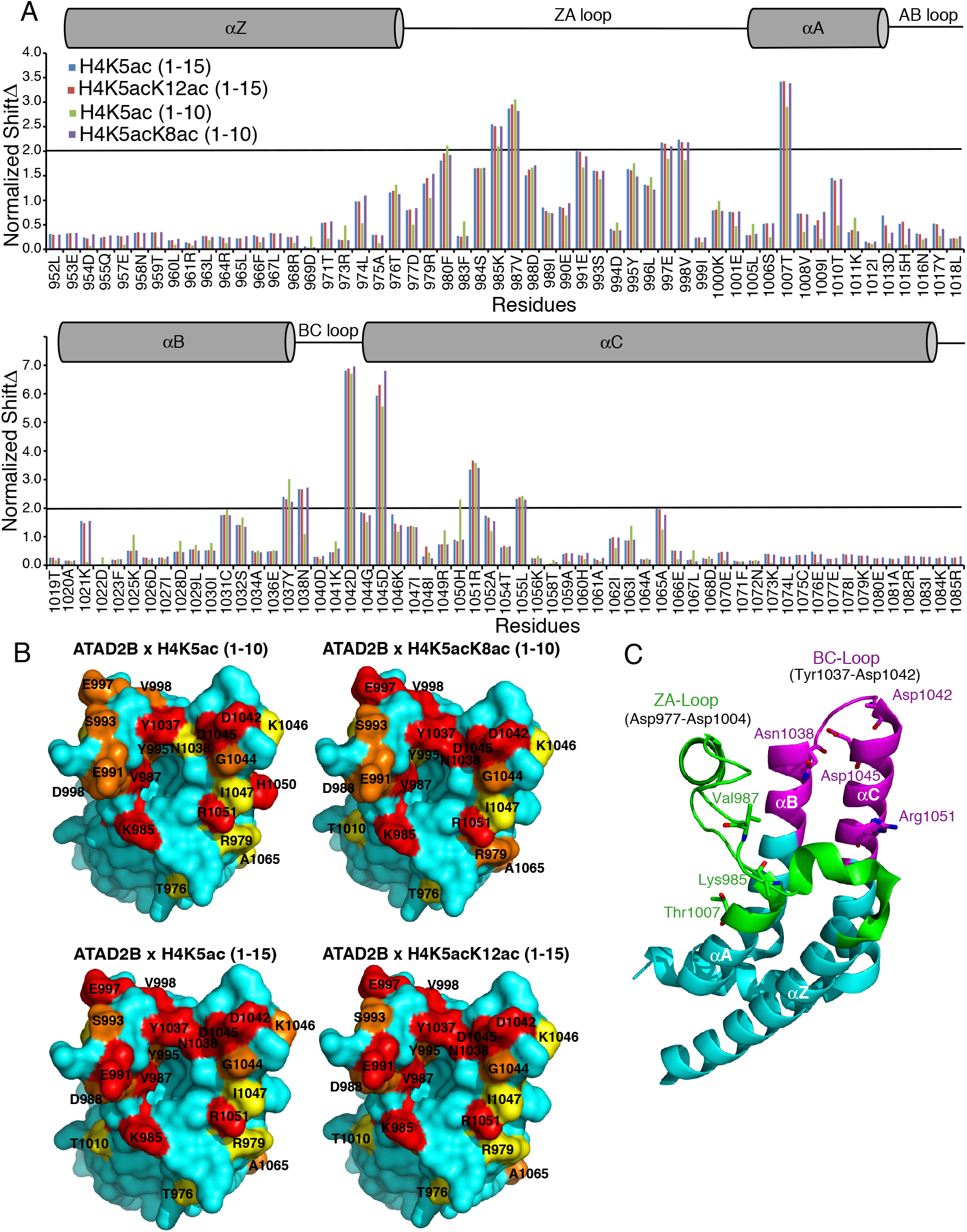
Mapping of the mono- and di-acetylated histone ligand binding interfaces on the ATAD2B bromodomain. (A) Histogram shows the normalized ^1^H-^15^N chemical shift changes observed in the backbone amides of the ATAD2B bromodomain upon addition of peptides H4K5ac (1-10) (green), H4K5acK8ac (1-10) (purple), H4K5ac (1-15) (blue) and H4K5acK12ac (1-15) (red) in a 1:5 (protein: peptide) molar ratio. The secondary structure elements for the bromodomain is represented above the histogram. A cylinder indicates an alpha-helical structure and a straight line depicts loop regions. (B) Chemical shift perturbations mapped onto the solvent-accessible surface representation of the apo ATAD2B bromodomain (PDB ID: 3LXJ) with the addition of H4K5ac(1-10), H4K5acK8ac(1-10), H4K5ac(1-15), H4K5acK12ac (1-15). CSPs that are 0.5, 1 and 2 standard deviations from the average shift changes are colored yellow, orange and red, respectively. (C) Cartoon representation of the ATAD2B bromodomain with the ZA loop region (residues 977-1004) colored in green and the extended region around the BC loop (residues 1031-1055) involved in chemical shift perturbations in magenta. The seven amino acids residues exhibiting the largest chemical shift changes are labeled in stick representation.

Consequently, mapping of these residues on the crystal structure of human ATAD2B bromodomain (PDB ID: 3LXJ) helped visualize the histone ligand interaction site (**Fig. 2B**). Overall, the residues showing large chemical shift perturbations upon addition of mono- and di-acetylated histone ligands are mainly found in two distinct regions of the ATAD2B bromodomain backbone. The first region spans residues 985-1007 which also overlaps with the variable ZA loop and short αAZ helix. The second region includes residues 1037-1055 that form the BC loop and the N-terminus of the αC helix region in the ATAD2B bromodomain structure (highlighted in green and magenta respectively, in **Fig. 2C)**. It is interesting to note that certain residues located outside of the canonical Kac binding pocket also showed large CSPs (above 2σ) (e.g. Thr 1007 and Arg 1051). We suspect this is most likely due to conformational changes occurring within these residues upon histone ligand binding.

### Coordination of mono-versus di-acetylated histone ligands

We also compared the differences in coordination of the mono- versus di-acetylated histone ligands (**Fig. 2A**). Overall, a larger number of differences in the chemical shift perturbation pattern was observed between the H4K5ac (1-10) and H4K5acK8ac (1-10) ligands than between the H4K5ac (1-15) and H4K5acK12ac (1-15) peptides. Notably, longer histone peptides increased the number of CSPs with a standard deviation greater than 2, but only minor differences were seen in the chemical shift perturbation pattern between the H4K5ac (1-15) and H4K5acK12ac (1-15) ligands. These CSPs occur in residues Arg 979, Phe 980 Val 987, Asp 988 in the ZA loop region, as well as Asp 1045 and Arg 1051 in the αC helix. However, since the mono-acetylated histone H4K5ac (1-15) peptide showed a ~3-fold increase in binding affinity over the H4K5acK12ac (1-15) peptide (as measured by ITC), the limited difference in the CSP pattern could indicate that the second Kac group is not coordinated directly by the ATAD2B bromodomain, but rather lies outside the binding pocket. On the other hand, there are several significant differences in the CSP pattern between the shorter histone H4K5ac (1-10) and H4K5acK8ac (1-10) ligands. CSP differences greater than 1σ were observed for residues Thr 1010, Lys 1021, Asn 1038, and Asp 1045. Residues Leu 974, Arg 979, Thr 1007, Tyr 1037, and Ala 1065 all demonstrated a CSP difference between 0.5-1.0σ. Thus, these residues displaying larger chemical shift patterns upon addition of the di-acetylated histone H4K5acK8ac (1-10) ligand are either due to conformational changes occurring in the ATAD2B bromodomain binding pocket upon complex formation, or due to the direct interaction of these residues with the second Kac group.

### Bromodomain binding pocket residues important for histone ligand recognition

Next, we sought to investigate the residues critical for the bromodomain-ligand interaction. In cloning the full-length ATAD2B cDNA for functional studies we discovered a splice variant in the ATAD2B mRNA that is created by an alternative 5’ splice donor site between exons 21 and 22 of ATAD2B that encode the bromodomain (**Fig. 3A and 3B**). This novel splicing event results in the exclusion of residues 988-992 (VDIEE) in the ZA loop (**Fig. 3C**) of the canonical ATAD2B bromodomain, generating a shorter variant (ATAD2B_short_). Interestingly, despite having an overall comparable gene structure to *ATAD2B*, we failed to detect any splicing events in the *ATAD2* bromodomain exons (**Supplementary Fig. 4A and 4B**). We used ITC assays to compare the ligand binding of the ATAD2B_short_ to the canonical ATAD2B bromodomain. In addition, we tested several conserved amino acids in this region via site-directed mutagenesis, including Tyr 995, Tyr 1037, and Asn 1038 residues. The locations of each of these amino acids are mapped onto the ATAD2B bromodomain binding pocket in **Fig. 3C**. For these binding assays, we used the five major ATAD2B bromodomain histone ligands identified from our ITC assays (**Table 2 and Supplementary Fig. 5**). Mutation of residues Tyr 995 and Tyr 1037 to alanine abolished binding to the mono- or di-acetylated histone ligands. In contrast, the ATAD2B N1038A mutation did not fully prevent histone ligand binding, as reduced binding activity for the major ATAD2B histone ligands was still observed (**Table 2 and Supplementary Fig. 5**). Interestingly, the ATAD2B_short_ bromodomain is unable to bind to any of the histone peptide ligands tested (**Table 2 and Supplementary Fig. 5)**. To verify that the loss of histone ligand binding was not due to protein misfolding after the mutation, we used circular dichroism (CD) to analyze the secondary structures of each of the tested ATAD2B bromodomains (**Table 3 and Fig. 3D**). Analysis of the ATAD2B mutants and the ATAD2B_short_ variant revealed no major alterations in the bromodomain three-dimensional structure. However, we further characterized the melting temperature (T_m_) of the ATAD2B bromodomain by a thermal shift assay and found that the ATAD2B_short_ variant has a significantly lower T_m_ when compared to the full-length bromodomain (**Fig. 3E**), presumably due to destabilization of the bromodomain fold. Overall, our data highlights the critical importance of the ZA loop in ligand recognition and suggests that alternative splicing could be a cellular mechanism regulating ATAD2B bromodomain ligand recognition.

**Figure 3.**
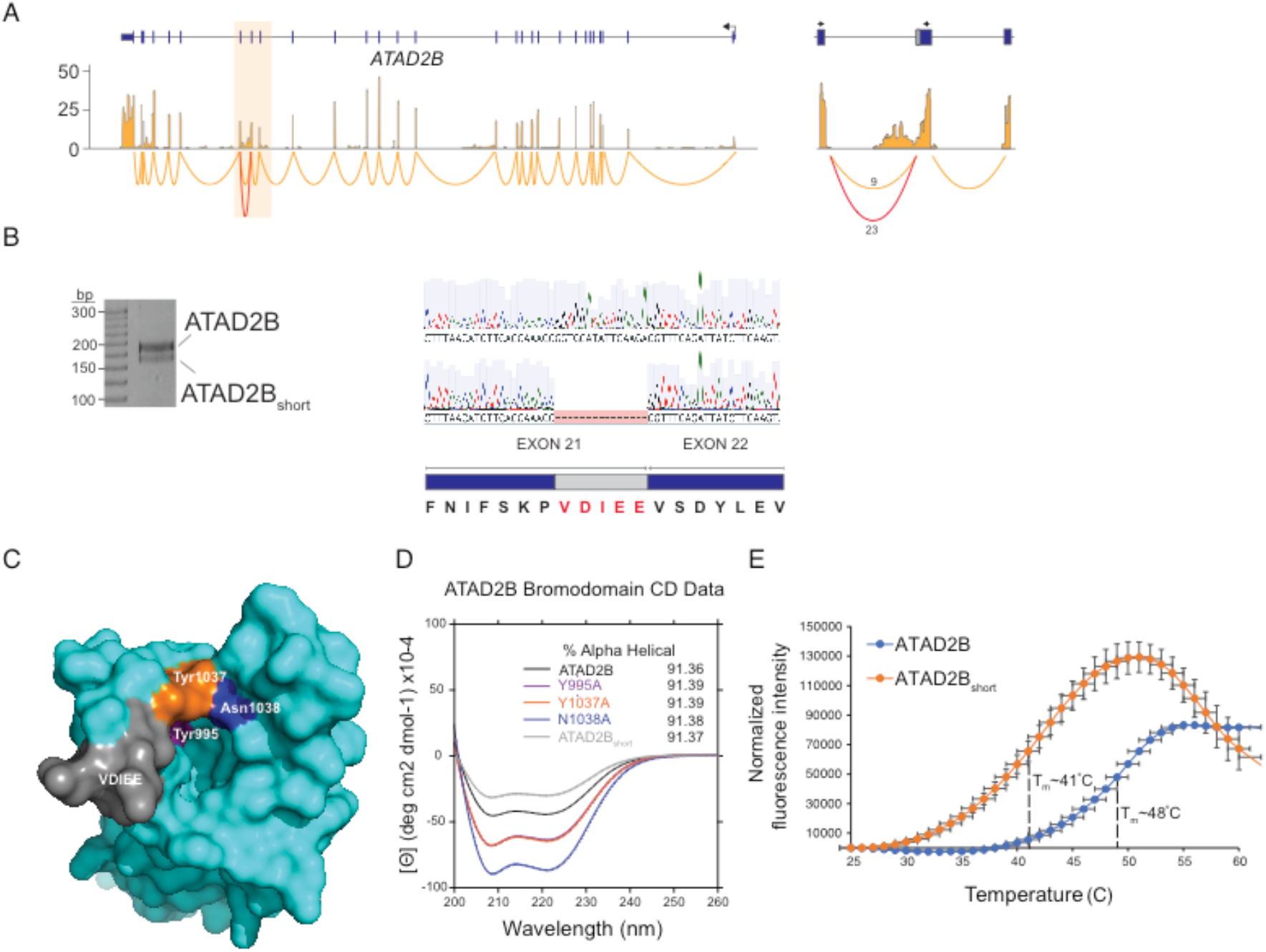
Analysis of targeted mutations and non-canonical splice variant in the ATAD2B bromodomain. (A) RNA-seq and sashimi plots showing expression and splice junctions across the *ATAD2B* gene. Normalized expression values are on the y-axis. The highlighted region indicates exons 20-22 that encode the *ATAD2B* bromodomain. The junctions for these exons are shown on the right where the number of reads spanning each junction is shown and the non-canonical splice variant is in red. (B) RT-PCR and Sanger sequencing showing both ATAD2B and ATAD2B_short_ isoforms. The amino acids excluded from ATAD2B_short_ as a result of alternative 5’ splice donor site are highlighted in red. (C) Surface representation of the apo ATAD2B bromodomain showing the location of specific point mutations introduced to the binding pocket by site-directed mutagenesis and the location of the residues excluded from the ATAD2B_short_ splice variant. (D) Circular dichroism spectra in the far-UV region of the ATAD2B, ATAD2B mutant, and ATAD2B_short_ bromodomains. The percent alpha-helical content for each protein is listed in the insert. (E) Thermal stability assay showing the ATAD2B bromodomain unfolding curve (blue) with a T_m_ = 48°C, while the ATAD2B_short_ bromodomain (orange) demonstrated a significantly lower T_m_ = 41°C.

**Table 2.** Dissociation constants of the wild-type and mutant ATAD2B bromodomain proteins with histone ligands as measured by ITC experiments. Dissociation values are micromolar (μM) unless otherwise noted.

| **ATAD2B bromodomain** | **H2AK5ac (1-12)** | **H4K5ac (1-10)** | **H4K12ac (4-17)** | **H4K5acK8 ac (1-10)** | **H4K5acK12 ac (1-15)** | **H4 unmodified (4-17)** | **Compound 38** |
| --- | --- | --- | --- | --- | --- | --- | --- |
| ATAD2B | $42.5 \pm 2.9$ | $48.0 \pm 4.7$ | $50.5 \pm 5.8$ | $28.1 \pm 1.6$ | $18.7 \pm 0.9$ | No binding | $160.6 \pm 12.3$ (nM) |
| N1038A | $1697.0 \pm 524.1$ | $720.5 \pm 4.1$ | No binding | $622.8 \pm 38.5$ | $411.7 \pm 39.1$ | No binding | --- |
| Y995A | No binding | No binding | No binding | No binding | No binding | No binding | --- |
| Y1037A | No binding | No binding | No binding | No binding | No binding | No binding | --- |
| ATAD2B<sub>short</sub> | No binding | No binding | No binding | No binding | No binding | No binding | No binding |

**Table 3.**
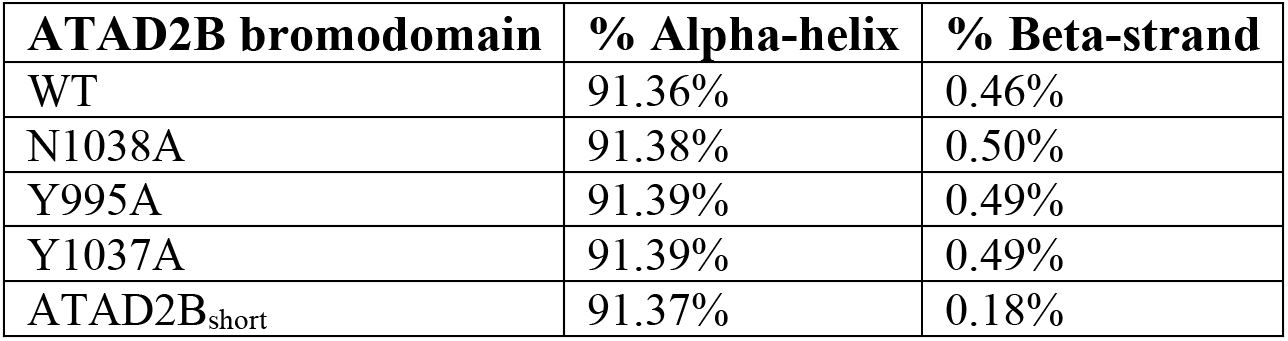
Analysis of the ATAD2B bromodomain wild-type and mutant proteins by circular dichroism spectroscopy.

**Figure 4.**
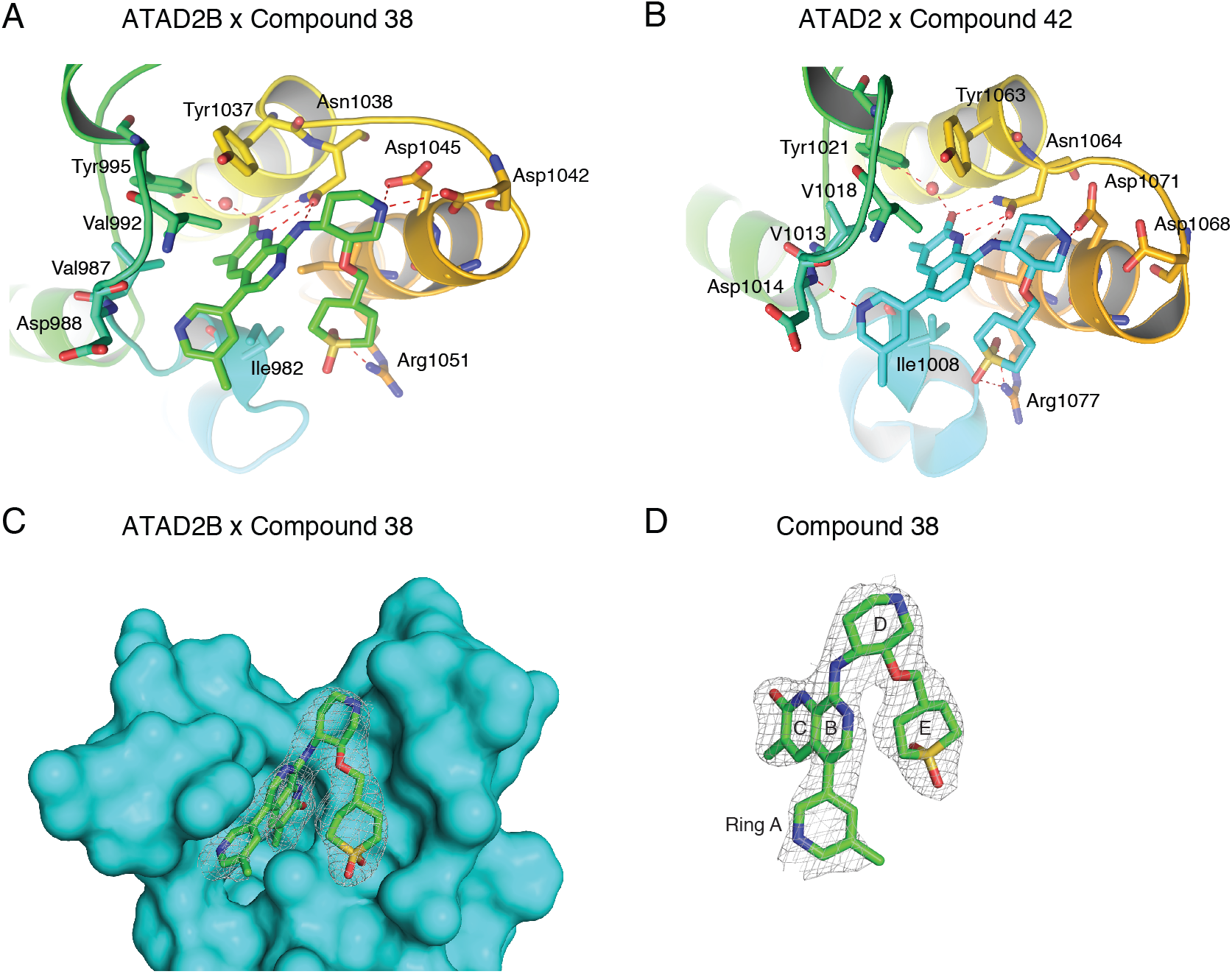
Coordination of bromodomain inhibitor molecules by the ATAD2 and ATAD2B bromodomains. (A) The ATAD2B bromodomain in complex with compound 38. (B) The ATAD2 bromodomain in complex with compound 42. Hydrogen bonds are indicated by a red dashed line. (C) The simulated annealing composite omit map (grey) representing the bound compound 38 (green) in complex with the ATAD2B bromodomain (cyan) contoured at 1σ. (D) Isolated image of the simulated annealing composite omit map (grey) around compound 38 (green) contoured at 1σ. The simulated annealing composite omit maps shown in (C) and (D) were calculated prior to building the ligand into the structure. Figures were generated with the PyMOL Molecular Graphics System, Version 2.3 Schrödinger, LLC.

### Recognition of the ATAD2 bromodomain inhibitor compound 38 (GSK C-38)

ATAD2 overexpression is strongly correlated with cancer progression^20–22^, driving the search for potent ATAD2 bromodomain inhibitors. Compound 38 (C-38) was developed by GlaxoSmithKline as the lead molecule possessing significant binding selectivity for the ATAD2 bromodomain over the BRD4 BD1 bromodomain (BET bromodomain subfamily II), with a binding affinity of 90 nM^17^. We tested the binding of C-38 against the ATAD2/B bromodomains by ITC and confirmed that this inhibitor interacts with the ATAD2 bromodomain with a *K*_D_ of 93.3 ± 2.8 nM, while the ATAD2B bromodomain binds with a *K*_D_ value of 160.6 ± 12.3 nM (**Table 1 and Supplementary Fig. 2S**). Furthermore, the ATAD2B_short_ VDIEE variant failed to interact with C-38 (**Table 2**).

To gain insights on why the binding affinity of the ATAD2B bromodomain for C-38 is almost 2-fold lower than the ATAD2 bromodomain^17^, we used molecular replacement with the apo ATAD2B bromodomain (PDB ID: 3LXJ) to solve the X-ray crystal structure of this compound in complex with the ATAD2B bromodomain. This structure provides the molecular details of the inhibitor interaction (**Table 4 and Fig. 4**) and demonstrates that coordination of C-38 occurs through a combination of hydrogen bonds and hydrophobic interactions (**Fig. 4A)**. The guanidinium group of Arg 1051 makes specific hydrogen bond contacts to the sulfone oxygen atoms of ring E in C-38, while the OD1 and OD2 carboxyl groups of Asp 1042 and Asp 1045 contact the piperidine nitrogen in ring D. The universally conserved Asn 1038 makes several hydrogen bonds to C-38. Asn 1038 binds the exocyclic amino group between the D and B rings, and forms H-bonds with both the nitrogen and oxygen atoms in the lactam ring C. The hydroxyl group of Tyr 995 also coordinates the oxygen atom of the lactam ring C through an ordered water molecule. Important hydrophobic interactions occur between C-38 and the side chains of the gatekeeper residue Ile 1048 and Ile 982 in the NIF shelf of the ATAD2B bromodomain (RVF shelf in ATAD2). Val 992, Val 987 and Tyr 1037 also contribute to the hydrophobic coordination of C-38 by forming part of the ATAD2B bromodomain binding pocket wall. The conserved Asn 1038 seems to play a large role, especially deeper in the binding pocket and is likely the driving force maneuvering the inhibitor into the bromodomain binding pocket in a favorable alignment. Of note, many of the bromodomain residues involved in the coordination of the C-38 inhibitor also make conserved contacts to the acetylated histone ligands (**Fig. 2 and Table 2**). Overall, our structural model indicates that coordination of C-38 by the ATAD2B bromodomain is highly conserved with the ATAD2 bromodomain. However, some key differences that may contribute to the change binding affinity are highlighted below.

**Table 4.** Summary of the data collection and refinement statistics for the ATAD2B bromodomain in complex with C-38.

| (collection on a single crystal) | ATAD2B / compound 38 |
| --- | --- |
| PDB code | 6VEO |
| **Data collection** |  |
| Space group | P6<sub>1</sub>22 |
| Cell dimensions |  |
| *a*, *b*, *c* (Å) | 76.358, 76.358, 102.312 |
| $\alpha$, $\beta$, $\gamma$ (°) | 90, 90, 120 |
| Resolution (Å) | 19.55 - 2.4 (2.486 - 2.4*) |
| R-pim (%) | 1.997 (17.07) |
| CC<sub>1/2</sub> | 1 (0.909) |
| Mean I/$\sigma$(I) | 21.51 (3.95) |
| Completeness (%) | 99.61 (100.0) |
| Redundancy | 2.0 (2.0) |
| **Refinement** |  |
| Resolution (Å) | 19.55-2.4 |
| No. unique reflections | 7332 (709) |
| R<sub>work</sub>/R<sub>free</sub> | 19.29 / 25.48 |
| No. atoms: |  |
| Protein | 1138 |
| Ligand/ion | 46 |
| Waters | 100 |
| Average B-factors (Å<sup>2</sup>) |  |
| Protein | 43.9 |
| Ions | 51.9 |
| Water | 47.8 |
| R.M.S deviations: |  |
| Bond lengths (Å) | 0.005 |
| Bond angles (°) | 0.55 |
| Ramachandran plot (%) |  |
| Favored | 97.76 |
| Allowed | 2.24 |
| Outliers | 0 |
| Clash score | 2.98 |

\*Highest resolution shell is shown in parenthesis.

## DISCUSSION

The ATAD2 bromodomain, which is 74.7% identical and 94.4% similar to the ATAD2B bromodomain, has been studied extensively due to its adverse association with overall survival rates for several solid tumors^21,23–26^. Previous studies have identified several histone-ligand interactions of the ATAD2 bromodomain, however little is known about the paralogous bromodomain found in ATAD2B^7,15,16^. Our dCypher peptide array data show that the ATAD2/B bromodomains predominantly select for acetyllysine modifications on histone H4; however, the ATAD2B bromodomain demonstrates a broader substrate specificity, not only binding to a larger number of histone H4 ligands, but also interacting with acetyllysine on histones H2A and H2A.X. The ATAD2B bromodomain is also able to recognize larger acyl modifications including propionyllysine and butyryllysine, but not crotonyllysine, similarly to the BRD9 family IV bromodomain^10^.

Prior to this study, Leachman *et al.*, 2010^12^ published one of the only sources of information on the ATAD2B protein. They studied the expression of ATAD2B in chicken embryos and human tumors with a polyclonal antibody raised against human ATAD2B. In 2012, the X-ray crystal structure of the ATAD2B (KIAA1240) bromodomain was deposited as an apo structure (PDB ID: 3LXJ) as part of a large-scale structural analysis of human bromodomains^7^. In the SPOT peptide array, ATAD2B demonstrated very strong interactions with acetylated histone peptides H2AK36ac, H2BK43ac, H2BK85ac, H3K56ac, H4K5ac, and H4K59ac^7^. It is clear from our experience determining the histone ligands of the BRPF1 bromodomain that high-throughput methods are important for identifying potential ligands, but more traditional biochemical and biophysical approaches are necessary to validate the interactions and measure the binding affinities^27^. It is not certain if acetylated lysines found outside of the N-terminal histone tail region (usually defined as the first 30 amino acids) would be sterically available for bromodomain recognition since they are located in the globular core domain of the histone proteins. Unfortunately, more specific follow up assays were not performed on the ATAD2B bromodomain to confirm these interactions.

Previous studies indicated that the ATAD2 bromodomain has a stronger binding preference for histone H4K12ac over H4K5ac, with reported affinities of 2.5 μM^14^ and 22 μM^16^, respectively. In contrast, our ITC binding data demonstrated that the ATAD2B bromodomain preferentially selects for the histone H4K5ac ligand over H4K12ac. Additionally a TR-FRET (Time-Resolved Fluorescence Energy Transfer) assay showed that the ATAD2 bromodomain recognizes multiple acetyllysine combinations on histone H4 including H4K5acK12ac (1-25) and H4K5acK8acK12acK16ac (1-25)^16^. We also found that the ATAD2B bromodomain recognizes several di-acetylated histone ligands including H4K5acK8ac, H4K5acK12ac, H4K5acK16ac, H4K8acK16ac, and H4K12acK16ac, albeit with weaker affinity. Both bromodomains appear to have overlapping ligand binding ability and recognize H4K5acK12ac^16^. Recognition of this di-acetylated histone H4K5acK12ac ligand was shown to be important for the biological function of the ATAD2 bromodomain, as this modification appears on newly synthesized histones following DNA replication^16,28^. The histone H4K5acK8ac modification has been linked to highly active gene promoters^29^. A previous study proposed that acetylation of histone H4 occurs progressively, starting at H4K16ac, then moving towards the N-terminus via K12, K8, and finally K5^30,31^. This model suggests that the H4K5acK8ac mark represents a hyper-acetylated state of histone H4^32^. Since our ITC data demonstrated a high binding affinity of the ATAD2B bromodomain with the H4K5acK8ac ligand, our results are the first to show a possible association of ATAD2B with hyperacetylated histone H4. Overall, our data indicate that the ATAD2/B bromodomains possess slightly different histone binding specificities despite being highly conserved paralogs.

The flexibility of the bromodomain reader module appears to be a key factor in the selection and recruitment of histone ligands. Using a molecular dynamics (MD) approach Langini *et al*., 2017^33^ simulated the ATAD2 bromodomain binding trajectories of the H4K5acK12ac (1–16) ligand with either a buried K5ac, K12ac, or just the bromodomain in the apo state^33^. They highlighted the role of Glu 1017, Tyr 1021, and Ile 1074 in ATAD2 (corresponding to Glu 991, Tyr 995, and Lys 1048 in ATAD2B), in the ZA loop that engage in “open” or “closed” conformational states of the loop. They also suggested that several negatively charged residues on the ZA loop could play a key role driving binding of the histone tail, which is composed of positively charged residues such as arginine and lysine. Our NMR results indicate significant chemical shift perturbations for residues Glu 991 (>1σ), Tyr 995 >1σ) and residue Glu 997 (~2σ) occur during titrations with mono- or di-acetylated H4 histone tail ligands, thus confirming the involvement of the bromodomain ZA loop in coordinating the histone ligands. The MD simulations of the ATAD2 bromodomain also detected molecular contacts between the histone ligand and a patch of negatively charged residues on the ZA loop^33^. These residues included Glu 1017 and Asp 1020, which correspond to residues Glu 991 and Asp 994 in the ATAD2B bromodomain. Previously, large conformational changes were observed in these regions of the ATAD2 bromodomain structure as reported by Poncet-Montange, *et al*., 2015^14^, however, our NMR data show that larger chemical shift perturbations occur in the BC loop than in the ZA loop region in the ATAD2B bromodomain. Moreover, many of the conserved residues involved in ligand coordination for the ATAD2 bromodomain, such as the Phe in the RVF shelf residues (ATAD2 1007-1009, ATAD2B 981-983) and the gatekeeper residue (Ile 1074/Ile 1048), did not demonstrate significant (>0.5σ) chemical shift perturbations upon ligand binding to ATAD2B. From these data, we hypothesize that Kac ligand recognition is initiated by structural changes in the flexible loop regions of the ATAD2B bromodomain that recruits the histone ligand to the bromodomain binding pocket resulting in specific hydrogen bond and hydrophobic interactions following ligand docking.

Interestingly, our previous NMR studies on the BRPF1 bromodomain (Family IV BRD) showed similar chemical shifts in both the ZA and BC loop regions^27^. Thus, inherent mobility in the ZA and BC loop regions appears to be a common feature of the bromodomain binding pockets, and they may retain unordered properties even when the ligand is bound. In **Fig. 2**, Glu 991 shows significant chemical shift changes upon the addition of nearly all histone ligands tested. Glu 991 corresponds to residue Asp 36 in the BRPF1 bromodomain, which also demonstrated one of the largest chemical shift perturbations upon ligand binding^27^. However, mutation of Asp 36 to Ala in the BRPF1 bromodomain did not significantly affect its histone binding activity, Therefore, we suspect that Glu 991 in ATAD2B may play an important role in the initial recruitment of the acetylated ligands into the bromodomain binding pocket. Overall, our data support the suggestion by Langini *et al*., 2017^33^ that dynamic motions in the binding pocket loops are likely important for the initial selection of the histone ligand, while ligand specificity is conferred by conserved amino acids that do not demonstrate large conformational changes upon ligand binding^33^. Thus, flexibility in the ZA and BC loop regions allow for rapid on/off rates of the histone ligand, and cooperative coordination of adjacent acetyllysine marks, but the universally conserved asparagine, the gatekeeper residue, and the NIF/RVF shelf motif in the ATAD2/B bromodomains drive histone ligand recognition at the molecular level.

Another important and interesting distinction between the ATAD2/B bromodomains is the discovery of a functionally distinct splice variant in the ATAD2B mRNA that is not found in the ATAD2 mRNA. Our data indicate that an alternative 5’ splice donor site of the ATAD2B bromodomain exons produces a bromodomain that lacks five amino acids in the ZA loop of the Kac binding pocket. Importantly, this variant does not interact with histone ligands, nor the small molecule inhibitor C-38. We performed directed RT-PCR splicing analysis and RNA-seq analysis of publicly available data^34^ to assess the expression patterns of ATAD2B bromodomain isoforms across a wide variety of cell types (**Supplementary Fig. 6**. Overall, we consistently detected comparable expression of the ATAD2B canonical and ATAD2B_short_ isoforms across different samples. There appears to be no direct correlation of ATAD2B_short_ isoform variant expression with breast cancer or across normal tissues.

Even though we failed to identify any evidence of alternative splicing in the ATAD2 bromodomain-encoding exons (**Supplementary Fig. 4**), such isoforms have been observed in other bromodomain-containing proteins. For example, BRPF1 bromodomain splice isoform A inserts six residues into the ZA-loop that effectively impairs the binding of tetra-acetylated histone H4 peptides and small molecule inhibitors^35^. Another splice variant observed in the NURF chromatin remodeling complex removes the C-terminal PHD finger and bromodomain and results in a spermatocyte arrest phenotype^36^. Additionally, a splice variant of p300 was described in a T-lymphoblastic cell line that generates a bromodomain lacking seven amino acid residues^37^. Thus, we hypothesize that alternative splicing may be a potential regulatory mechanism controlling ATAD2B activity *in vivo*.

Our ITC binding data on the interaction of the ATAD2B bromodomain with the GlaxoSmithKline ATAD2 bromodomain inhibitor C-38 (**Table 1 and Supplementary Fig. 2S**) is in-line with the previously published observation that C-38 binds well to both the ATAD2/B bromodomains in the nanomolar range^17^. In addition, we solved the structure of the ATAD2B bromodomain with C-38 and identified molecular contacts that are important for inhibitor coordination. The X-ray crystal structure of the ATAD2B bromodomain in complex with C-38 (PDB ID: 6VEO) is highly conserved with the structure of the ATAD2 bromodomain in complex with compound 42 (C-42, PDB ID: 5A83). C-42 was also developed by GlaxoSmithKline as a derivative of C-38 with increased cell membrane permeability^17^. C-38 differs from C-42 only by the presence of one nitrogen atom in the lactam ring B, which is a carbon in the latter. The coordination of C-42 in the ATAD2 bromodomain is nearly identical to the coordination of C-38 by the ATAD2B bromodomain (compare **Fig. 4A and 4B, and Supplementary Fig. 7)**. The hydrogen-bonding pattern is highly conserved, with important H-bonds formed to the universally conserved asparagine, Asn 1038 and Asn 1064 in the ATAD2/B bromodomains, respectively. In the ATAD2-C-42 structure there are also H-bonds to the backbone nitrogen of Asp 1014, and the side chains of Asp 1071 and Arg 1077.

Notably, there are slight differences in the H-bond coordination of C-38 and C-42 between the ATAD2B and ATAD2 bromodomains (**Fig. 4A-B**). For instance, the backbone nitrogen of Asp 1014 in ATAD2 makes an H-bond contact with the nitrogen atom of pyridine ring A in C-42. In ATAD2B the C-38 ligand is positioned closer to Asp 1042 and 1045, weakening the interaction between Asp 988 and the pyridine ring nitrogen, and loss of this contact is likely related to the lower affinity for C-38 by the ATAD2B bromodomain. The RVF shelf motif in the ATAD2 bromodomain is also the main region of the Kac binding pocket that differs between the ATAD2/B bromodomains, which is a NIF shelf motif in ATAD2B.

Overall the ATAD2/B bromodomain Kac binding pockets are very highly conserved and finding inhibitors that will differentiate between these two bromodomains will be challenging. Even so, a notable success was the discovery of BAY-850, an isoform-specific small molecule shown to selectively target only the ATAD2 bromodomain^38^. BAY-850 appears to have a unique mode of action and causes dimerization of the ATAD2 bromodomain to inhibit its interaction with chromatin. Thus, the optimization of new inhibitors to take better advantage of the small differences between the ATAD2/B bromodomains could successfully improve their binding specificity of bromodomain inhibitor compounds. Indeed, one region with significant differences between the ATAD2/B bromodomains is at the RVF shelf motif found in ATAD2 (residues Arg 1007, Val 1008 and Phe 1009), which forms an NIF shelf in the ATAD2B bromodomain (residues Asn 981, Ile 982 and Phe 983) (**Fig. 4A**).

## CONCLUSIONS

Collectively, the results from this study describe the histone ligands of the ATAD2B bromodomain and outline many important features of the binding pocket that contribute to ligand selection and specificity. This information will be valuable for the development of selective ATAD2B bromodomain inhibitors or tool compounds that can be used to further investigate the biological function of the poorly characterized ATAD2B protein. We discovered that the ATAD2B bromodomain is a di-acetyllysine reader module, however, the significance of histone H4K5acK8ac ligand binding has yet to be determined. Noteably, dysregulation of histone acetylation patterns has been linked to cancer development^39^, and certain Kac modifications have been shown to increase and/or decrease in different cancer subtypes^40^. Linking the recognition of histone H4K5acK8ac to particular cellular outcomes will be essential for understanding the unique role of the ATAD2B bromodomain, and how it functions independently from ATAD2. Our NMR chemical shift perturbation experiments identified several amino acids that differed between the coordination of mono- and di-acetylated histone ligands. It is interesting to note that most of the differences between the chemical shift perturbation pattern in the coordination of H4K5ac (1-15) and H4K5acK12ac (1-15) occur in residues that are also involved in hydrogen bond contacts with the C-38 inhibitor. Additionally, differences in the chemical shift perturbation pattern of key residues provide new insights as to why this bromodomain preferentially selects for histone ligands with multiple Kac modifications.

Thus, it will be essential to solving the structure of the ATAD2B bromodomain with di-acetylated histones to discover the molecular details of the binding mechanism. However, our results demonstrate that although the ATAD2/B bromodomain modules are highly conserved paralogs, they preferentially recognize different subsets of acetylated histone ligands and appear to be regulated via different cellular mechanisms. This suggests that these two bromodomain-containing proteins are involved in divergent biological pathways, and may contribute to cancer development in different ways. It is also important to note that while the ITC and NMR experiments were robust, our biochemical and biophysical analysis may not be directly translated to the cell, where the *in vivo* abundance of the H4K5acK8ac and H4K5acK12ac modifications may be relatively small compared to either the H4K5ac or H4K12ac ligands, which also bind strongly to the ATAD2B bromodomain. Thus, further *in vivo* experimentation will be necessary to fully appreciate the biological significance of these interactions. In conclusion, we describe here the novel histone binding specificity of the ATAD2B bromodomain. Our study outlines the important residues involved in mono- and di-acetylated histone ligand recognition, as well as in coordination of the small molecule bromodomain inhibitor C-38. Our study also revealed that alternative splicing may be a unique molecular mechanism regulating the binding activity of the ATAD2B bromodomain. These data provide insight into how they may be specifically inhibited to further advance our understanding of differences in the roles of the ATAD2/B bromodomains.

## METHODS

### Plasmid Construction

Human ATAD2/B cDNA (UniProt codes Q9ULIO and Q6PL18) was a gift from Nicole Burgess-Brown (Addgene plasmid # 38916 and 39046)^7^. The ATAD2 bromodomain region (residues 981-1108) and the ATAD2B bromodomain region (residues 953-1085) were PCR-amplified and cloned into the pDEST15 vector encoding an N-terminus GST tag using the Gateway Cloning Technology (Invitrogen)^27^. ATAD2B bromodomain mutants N1038A, Y995A, Y1037A and the DIEEV deletion (residues 988-992) were generated using QuikChange^®^ mutagenesis (Agilent Technologies)^41^. The DNA sequence of each was verified (University of Vermont Cancer Center Advanced Genome Technologies Core) and then transformed into *Escherichia coli* Rosetta 2(DE3)pLysS competent cells (Novagen).

### Protein Expression and Purification

*E. coli* cells containing the GST-tagged ATAD2B wild-type or mutant bromodomain were grown in 4 L Terrific broth (TB) or ^15^NH_4_Cl-supplemented or ^15^NH_4_Cl/^13^C_6_ D-glucose-supplemented minimal media at 37°C. The culture temperature was dropped to 20°C for one hour when the OD_600_ reached 1. The culture was then induced by the addition of 0.25 mM isopropyl β-D-1-thiogalactopyranoside (IPTG), and the cells were harvested by pelleting after 16 h of incubation at 20°C with shaking at 225 rev min^−1^. The bacterial pellet was re-suspended in 100 mL lysis buffer (50 mM Tris–HCl pH 7.5, 500 mM NaCl, 0.05% Nonidet P-40, 1 mM DTT) containing 1 mL lysozyme and the cells were lysed by sonication. The cell lysate was cleared by centrifugation (10,000 rev min^−1^ for 20 min). The supernatant was added to 15 mL of glutathione agarose resin (which binds up to 40 mg of purified recombinant GST protein per milliliter of resin) (Thermo Scientific) and incubated on ice (4°C) while agitating for 120 minutes. After incubation, the suspension was centrifuged for 5 min at 500 x g to collect the beads. The collected beads were poured into a 25 mL Econo-Column Chromatography Column (Bio-Rad) and washed with four column volumes of wash buffer (20 mM Tris–HCl pH 8.0, 500 mM NaCl, 1 mM DTT). The GST tag was cleaved overnight at 4°C by the addition of PreScission Protease (~100 μL at 76 mg/mL) (GE Healthcare) and the eluted ATAD2B bromodomain protein was concentrated to a total volume of approximately 3 mL. For Isothermal Titration Calorimetry (ITC) or circular dichroism (CD) experiments protein was concentrated and dialyzed into buffers consisting of 20 mM NaH_2_PO_4_ pH 7.0 and 150 mM NaCl, or 50 mM NaH_2_PO_4_ pH 7.0 and 50 mM NaCl, respectively. For NMR experiments the uniformly ^15^N labeled and the ^15^N, ^13^C double-labeled ATAD2B bromodomain was buffer exchanged into 20 mM Tris-HCl pH 6.8, 150 mM NaCl, 10 mM DTT, and 10% D_2_O. For X-ray crystallography the protein was further purified using fast-protein liquid chromatography on an AKTA Purifier UPC 10 (GE Healthcare) over a HiPrep 16/60 Sephacryl S-100 high-resolution gel filtration column (GE Healthcare) equilibrated with crystallography buffer (25 mM HEPES pH 7.5, 150 mM NaCl and 1 mM DTT). Eluted fractions corresponding to the ATAD2B bromodomain were pooled and concentrated to 30 mg mL^−1^ at 4°C. The protein concentration was determined using the Pierce BCA Protein Assay Kit (Thermo Scientific) and was calculated from the absorption at 550 nm and the ATAD2B bromodomain extinction coefficient of 4470 *M* ^−1^ cm^−1^. The purity of the ATAD2B bromodomain was verified by SDS–PAGE gels stained with GelCode Blue Safe protein stain (Thermo Scientific).

### Histone Peptide and Bromodomain Inhibitor Synthesis

Biotinylated histone peptides for the dCypher assays were from EpiCypher (**Resources Table A**). Histone peptides for other studies were synthesized by the Peptide Core Facility at the University of Colorado Denver or by GenScript with a free N-terminus and an amidated C-terminus. The peptides, supplied as the TFA salt, were purified to greater than 98% pure via HPLC, and their chemical identities confirmed by mass spectroscopy (for sequence information see **Table 1**). Compound 38 (C-38) or 8-(((3R,4R)-3-((1,1-Dioxidotetrahydro-2H-thiopyran-4-yl)-methoxy)piperidin-4-yl)amino)-3-methyl-5-(5-methylpyridin-3-yl)-1,7-naphthyridin-2(1H)-one, is a potent ATAD2 bromodomain inhibitor developed by GlaxoSmithKline in 2015^17^. C-38 was synthesized by Dr. Jay Bradner’s laboratory according to the previously published procedure and isolated as the TFA salt^17^.

### dCypher binding assays

Two phases of dCypher^**®**^testing on the AlphaScreen^**®**^platform were performed as previously described^42,43^. In Phase A, the GST-tagged ATAD2/B bromodomains were titrated to positive and negative control biotinylated peptides to determine binding curves (including level of sensitivity, signal saturation, and signal over background: **Supplementary Fig. 1A-B**). In Phase B, a single optimal GST-bromodomain concentration was used to probe the complete biotinylated peptide panel (**Resources Table A**). For each phase, 5 μL of the GST-tagged ATAD2/B bromodomain (4 nM and 500 pM, respectively) was incubated with 5 μL of biotinylated peptides at 100 nM for 30 min at room temperature in the epigenetics assay buffer (PerkinElmer AL008; diluted to 1x and supplemented with 1mM TCEP) in a 384-well plate. A mixture of 10 μL of 2.5 μg/mL glutathione acceptor beads (PerkinElmer) and 5 μg/mL streptavidin donor beads (PerkinElmer) was prepared in peptide assay buffer and added to each well. The plate was incubated for 60 min at room temperature in subdued lighting, and the AlphaLISA signal measured on a PerkinElmer 2104 EnVision (680-nm laser excitation, 570-nm emission filter ± 50-nm bandwidth). Each binding interaction was performed in duplicate. The dCypher assay signal was plotted separately for the ATAD2 bromodomain and the ATAD2B bromodomain array data, where the data was scaled by the strongest average fluorescent intensity signals for each bromodomain.

### Isothermal Titration Calorimetry

ITC experiments were carried out at 5°C with a MicroCal iTC200 instrument (GE Healthcare). Each of the ATAD2B bromodomain proteins and histone peptide samples were dialyzed into ITC buffer containing 20 mM NaH_2_PO_4_ pH 7.0 and 150 mM NaCl. Calorimetric titration was performed by titrating each histone tail ligand (5 mM) into 0.2 mM of the ATAD2B bromodomain protein in the sample cell in a series of 19 individual injections of 2 μL, at time intervals of 150 seconds with a stir speed of 750 rpm. These were preceded by a preliminary injection of 0.5 μL of the 5 mM peptide sample, which was excluded from the data integration and calculation of K_D_s. To determine the heat of dilution of the titrant peptides in the experimental buffer, control experiments were conducted under identical conditions. As part of data analysis, this was deducted from the experimental data. The obtained change-in-heat peaks were then analyzed by the Origin 7.0 program (OriginLab Corporation) and used to calculate binding affinities. Experiments in which binding occurred were performed in triplicate, while non-binding experiments were performed in duplicate.

### NMR Spectroscopy

All chemical shift perturbation experiments were conducted using uniformly ^15^N-labeled ATAD2B bromodomain prepared at 0.500 mM in buffer containing 20 mM Tris-HCl pH 6.8, 150 mM NaCl, 10 mM DTT and 10% D_2_O. 35 μL titration mixtures of the protein and peptide were made at concentration ratios of 1:0, 1:0.25, 1:0.5, 1:1.1, 1:2.5, and 1:5 for the unlabeled histone tail peptides containing specific acetyllysine modifications. 2D ^15^N HSQC (heteronuclear single quantum coherence) experiments for all samples were run at 25°C on a 600 MHz Bruker AVANCE III spectrometer equipped with a *z*-gradient 1.7 mm TCI probe at the National Magnetic Resonance Facility at Madison (NMRFAM). The NMR data were collected with ^1^H and ^15^N radio-frequency pulses applied at 4.745 parts per million (ppm) and 116 ppm, respectively. 1024 × 128 complex data points with spectral widths of 16 ppm and 30 ppm, respectively, were collected along the ^1^H and ^15^N dimensions, with 32 scans per FID and an inter-scan delay of 1.0 sec, resulting in a total experimental time of about 160 min for each HSQC spectrum. Transformed spectra at each titration point were overlaid and analyzed to characterize the relevant residues affected by the interaction between the ATAD2B bromodomain and the histone peptides. Normalized chemical shift changes were calculated as described in^44^ using the equation:

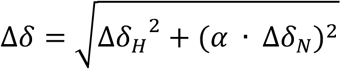

where α = 0.14 for all residues except glycine, which has α = 0.2, and Δδ_H_ and Δδ_N_, respectively, are the changes in proton and nitrogen chemical shift in ppm. To map the chemical shift perturbations onto the ATAD2B bromodomain surface, we calculated the standard deviation (σ) of the Euclidean chemical shift change over all residues of all bromodomain: peptide combinations. Residues whose chemical shift change was greater than 0.5σ were colored yellow, while residues with chemical shift changes greater than 1σ and 2σ were colored orange and red, respectively.

To obtain the backbone resonance assignments, the ^15^N, ^13^C double-labeled ATAD2B bromodomain was prepared at about 1 mM in buffer containing 20 mM Tris-HCl pH 6.8, 150 mM NaCl, 10 mM DTT and 10% D_2_O. ADAPT-NMR (Assignment-directed Data collection Algorithm utilizing a Probabilistic Toolkit in NMR) was used to optimize simultaneous fast data collection and automated NMR assignment of the HNCO, HN(CA)CB, HNCA, HN(CO)CA, HN(CA)CO, CBCA(CO)NH, and C(CCO)NH spectra by achieving reduced dimensionality (2D)^45^. These experiments were run at 25°C on a 600 MHz Agilent VNMRS spectrometer equipped with a 5 mm Z-axis pulsed-field gradient triple resonance cold probe. The NMR data for all experiments were collected with the universal carrier position of ^1^H, ^15^N, ^13^C^a^ (shaped pulse), ^13^C^aliphatic^ (^13^C^a/b^ or ^13^C^b^, shaped pulse), ^13^C’ (shaped pulse) applied at 4.76 ppm (H_2_O frequency), 118 ppm, 56 ppm, 45 ppm, and 176 ppm respectively. 1024, 32, 64, 64, and 64 complex data points with spectral widths of 16 ppm, 36 ppm, 32 ppm, 70 ppm, and 22 ppm, respectively, were collected along the ^1^H, ^15^N, ^13^C^a^, ^13^C^aliphatic^ and ^13^C’ dimensions. Processing and analysis for all data were conducted automatically with ADAPT-NMR^45,46^. After one day of the ADAPT-NMR run, an 84.7% assignment level was achieved. To generate visualized assignment labels on the 2D ^1^H-^15^N HSQC spectra, PINE-SPARKY and NMRFAM-SPARKY were used^47–49^.

### Circular Dichroism Spectroscopy

Circular Dichroism (CD) spectra were recorded on a JASCO J-815 CD Spectrometer (JASCO, Japan) at 25°C in a 1.6 cm cell. ATAD2B bromodomain wild type (WT) or mutant proteins were dialyzed in 50 mM NaPO_4_ pH 7.0 and 50 mM NaCl buffer and diluted to between 0.1 μM and 0.5 μM in concentration. CD spectra were measured from 199–260 nm. Two spectra were measured and averaged for each mutant bromodomain protein sample and the wild-type protein. Spectra were analyzed using the K2D3 structure prediction software to determine the percent α-helical and β-sheet content^50^.

### X-Ray Crystallography

The purified ATAD2B bromodomain protein (1.9 mM) was mixed with C-38 (4 mM) in a 1.5 mL microcentrifuge tube. Crystallization screens were set up using the sitting-drop method in 96-well VDX plates (Hampton Research), with drops consisting of 1 μL protein:peptide mixture plus 1 μL reservoir solution and a reservoir volume of 100 μL. The ATAD2B bromodomain was co-crystallized with the C-38 inhibitor in Index Screen condition No. 66 [0.2 M ammonium sulfate, 0.1 M Bis-Tris pH 5.5, 25% (w/v) polyethylene glycol 3,350] at 4°C. The C-38 crystals from Index Screen condition No. 66 were reproduced in hanging drops (1 μL protein solution plus 1 μL mother liquor) using 24-well VDX plates (Hampton Research) containing 500 μL mother liquor in the reservoir. The crystals were mounted directly from the drop using the CrystalCap SPINE HT system with 100 μM nylon loops (Hampton Research) with the notches broken at 18 mm. The crystals were cryoprotected by incubating the crystal in the cryoloop over 500 μL of a 40% methanol in water solution inside the cryovial where the liquid nitrogen escape holes were blocked with clay. After a 2 min incubation period, the crystal was flash-frozen directly in the liquid nitrogen stream at 100K on the beamline much as described in Farley *et al.,* 2014^51^. Data for the C-38 crystal was collected at the Center for X-ray Crystallography at the University of Vermont on a Bruker D8 Kappa Quest generator (Bruker-AXS Inc., Madison, WI, USA) equipped with a Photon 100 CMOS detector and a Cryostream 700 from Oxford Cryosystems. The diffraction data were processed using the Proteum3 suite (Bruker-AXS Inc., Madison, WI, USA). The structure was solved by molecular replacement using Phaser with the apo ATAD2B bromodomain structure (PDB ID: 3LXJ) as the starting model^52^. The model was built using COOT to add the C-38 inhibitor, and PHENIX and COOT were used for iterative rounds of refinement, density modification and model building^53,54^. The final structure at 2.4 Å resolution was validated using MolProbity and Polygon^55,56^.

### Thermal Shift Assay

The thermofluor assay or the thermal shift assay is often used for rapid determination of protein thermal stability in different *in vitro* conditions by monitoring the unfolding of the protein at increasing temperatures. The melting point (T_m_) of the wild-type or mutant ATAD2B bromodomain was calculated from the increase in the SYPRO orange dye fluorescence intensity. It was performed using a 96-well clear plastic plate using an Eppendorf Mastercycler Realplex2 qPCR machine. Triplicates of 100 μL of 5 μM ATAD2B WT and the ATAD2B DIEEV mutant were pre-mixed with 5 μM SYPRO orange dye in 20 mM sodium phosphate, 150 mM NaCl and pH 7.0 for T_m_ calculation. Wells containing only buffer with 5 μM dye were used for baseline correction. The plate was sealed with an optically clear adhesive tape to prevent sample loss during heating. The temperature was gradually ramped up from 25-95°C while monitoring the change in fluorescence intensity of the dye. GraphPad Prism software was used to analyze the data and calculate the melting point of the protein sample by applying nonlinear regression using the Melting Boltzmann equation:

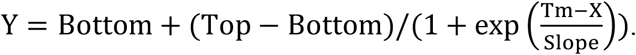

### RT-PCR analysis

Splicing analysis: Direct-zol RNA mini-prep kit (Zymo Research) was used to isolate RNA in Trizol reagent (Thermo Scientific) and cDNA synthesized using 1 μg of total RNA with Invitrogen™ SuperScript II Reverse Transcriptase and oligo dT primers (Thermo Scientific) according to the manufacturer’s instructions. PCR products were resolved on a 24% TBE gel and visualized using ethidium bromide or cloned into pGEM-T (Promega) and Sanger sequenced. Primer Sequences: ATAD2B_EXON_F: TGACCTCATCTGTAGCAATGCT and ATAD2B_EXON_R: TCAGCTGCAATGATAGCATGTGCA

### RNA-seq analysis

RNA-seq alignment files were obtained from the Cancer Cell-Line Encylopedia^34^. Reads spanning the splice junctions between exon 21 and exon 22 of ATAD2 and ATAD2B were quantified using seqsetvis^57^. Splicing patterns for *ATAD2* and *ATAD2B* were visualized using Gviz^58^, and splice junctions with counts less than 5 were removed for sashimi plots.

## Supporting information

Resource Table A

## ABBREVIATIONS

ATAD2: ATPase family AAA^+^ domain containing 2
ATAD2B: ATPase family AAA^+^ domain containing 2B
BRD: bromodomain
BRPF1/3: Bromodomain and PHD finger containing protein 1/3
CD: circular dichroism
GST: glutathione-S-transferase
HSQC: heteronuclear single quantum coherence
ITC: isothermal titration calorimetry
NMR: nuclear magnetic resonance
PHD: plant homeodomain
PTMs: post-translational modifications

## ACKNOWLEDGMENTS

We are especially thankful to Brian E. Eckenroth in the Department of Microbiology and Molecular Genetics at the University of Vermont for his assistance with crystallographic data collection, processing, and analysis of the ATAD2B bromodomain structure in complex with C-38. We are very grateful to Dennis L. Buckley and James E. Bradner at the Dana-Farber Cancer Institute/Harvard Medical School in Boston, MA for re-synthesizing and sharing compound 38 with us for use in this study. We also appreciate Kara McGuire’s and Juliet Obi’s assistance with the NMR chemical shift perturbation analysis and crystal growth, and Lauren Dunn’s help with cloning the ATAD2B VDIEE short construct, respectively. This study was supported by National Institute of Health grants: R01GM129338 to KCG and SF; R44GM116584 and R44GM117683 to EpiCypher; and R15GM104865 to KCG. JTL was the recipient of an ACPHS graduate research assistantship from 2016-2017. This study made use of the National Magnetic Resonance Facility at Madison, which is supported by NIH grant P41GM103399 (NIGMS). NMRFAM equipment was purchased with funds from the University of Wisconsin-Madison, the NIH P41GM103399, S10RR02781, S10RR08438, S10RR023438, S10RR025062, S10RR029220), the NSF (DMB-8415048, OIA-9977486, BIR-9214394), and the USDA. Crystal growth, screening, and initial data collection were carried out at the Center for X-ray Crystallography at the University of Vermont, which is supported by the National Institutes of Health Grants P01CA098993 and R01CA52040, awarded to Sylvie Doublié by the National Cancer Institute. We also thank Dr. Matthew Liptak and Amanda Roffman in the Department of Chemistry at the University of Vermont for their advice and assistance setting up circular dichroism spectroscopy experiments. DNA sequencing was performed at the University of Vermont Cancer Center Advanced Genome Technologies Core. The structure of the ATAD2B bromodomain in complex with compound 38 was deposited into the Protein Data Bank (PDB ID: 6VEO). GM/CA@APS has been funded in whole or in part with Federal funds from the National Cancer Institute (ACB-12002) and the National Institute of General Medical Sciences (AGM-12006). This research used resources of the Advanced Photon Source, a U.S. Department of Energy (DOE) Office of Science User Facility operated for the DOE Office of Science by Argonne National Laboratory under Contract No. DE-AC02-06CH11357.

## AUTHOR CONTRIBUTIONS

KCG, SF, JTL, and MP conceived and designed the experiments. M.R.M. and M.-C.K. performed the dCypher array and analyzed the data. JTL, MYL, JCG, MP, SB, CME, and SC performed the *in vitro* biochemical experiments with recombinant proteins and analyzed data. MT and GC collected the NMR data, and MP and KCG analyzed the data. DLB and JEB were responsible for the synthesis and validation of compound 38. BEE and KCG solved the crystal structure. KM, AD, CG, and SF carried out the genomic analysis, PCR, and cellular analysis of the ATAD2B splice variant. All authors contributed to the writing of the manuscript, reviewed the results, and approved the final version of the manuscript.

## COMPETING INTERESTS

EpiCypher (M.R.M. and M.-C.K.) is a commercial developer of the dCypher peptide-binding platform used in this study.

**Supplementary Tables 1 & 2**. Detection of modified histone ligands recognized by the GST-ATAD2/B bromodomains using the *dCypher* assay.

**Resources Table A**. Contains the specific identity of all histone peptides in the *dCypher* assays.

## Notes

### Summary of Updates

Significant new data was added to the current manuscript draft including, results from a histone peptide array screen of the ATAD2B bromodomain with nearly 300 unique PTM histone peptides, data identifying and characterizing an ATAD2B splice variant that abolishes histone and inhibitor binding, and inclusion of a new structure of the ATAD2B bromodomain in complex with a small molecule inhibitor with improved electron density. We also expanded the ITC binding data and reanalyzed the NMR titration data.

