## supplemental figures for "Structural insights into the recognition of mono- and di-acetylated histones by the ATAD2B bromodomain"

**Supplementary Figure 1. Characterization of the ATAD2/B bromodomain binding curves for the dCypher assay.** (A) The GST-tagged ATAD2 bromodomain (residues 981-1108) and (B) ATAD2B bromodomain (residues 953-1085) were titrated to control biotinylated histone peptides and binding curves determined on AlphaScreen® platform (i.e. dCypher® Phase A: see Methods). X-axes are log(protein concentration (M)) at constant peptide concentration (100 nM); Y-axes are AlphaScreen counts, representing relative strength of binding (n = 2; error bars are S.D.).

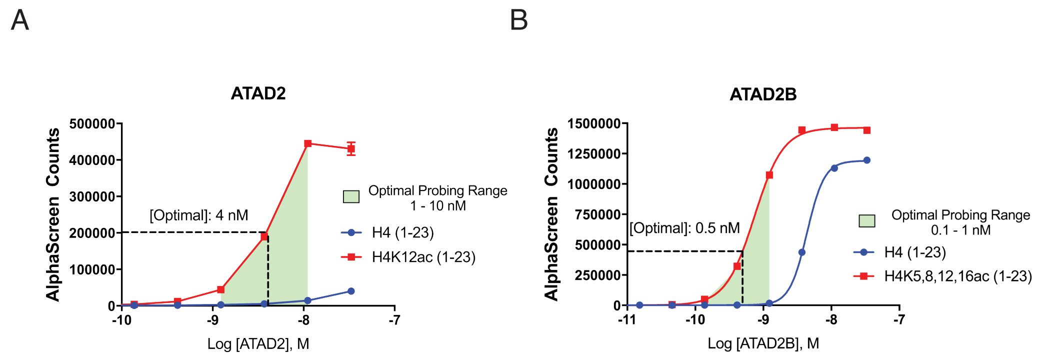

**Supplementary Figure 2. ITC measurements of the interaction between acetylated histone ligands and the ATAD2B bromodomain.** (A-S) Exothermic ITC enthalpy plots for the binding of the ATAD2B bromodomain to unmodified, mono- and di-acetylated histone ligands, as well as the ATAD2 bromodomain inhibitor compound 38. The calculated binding constants are indicated.

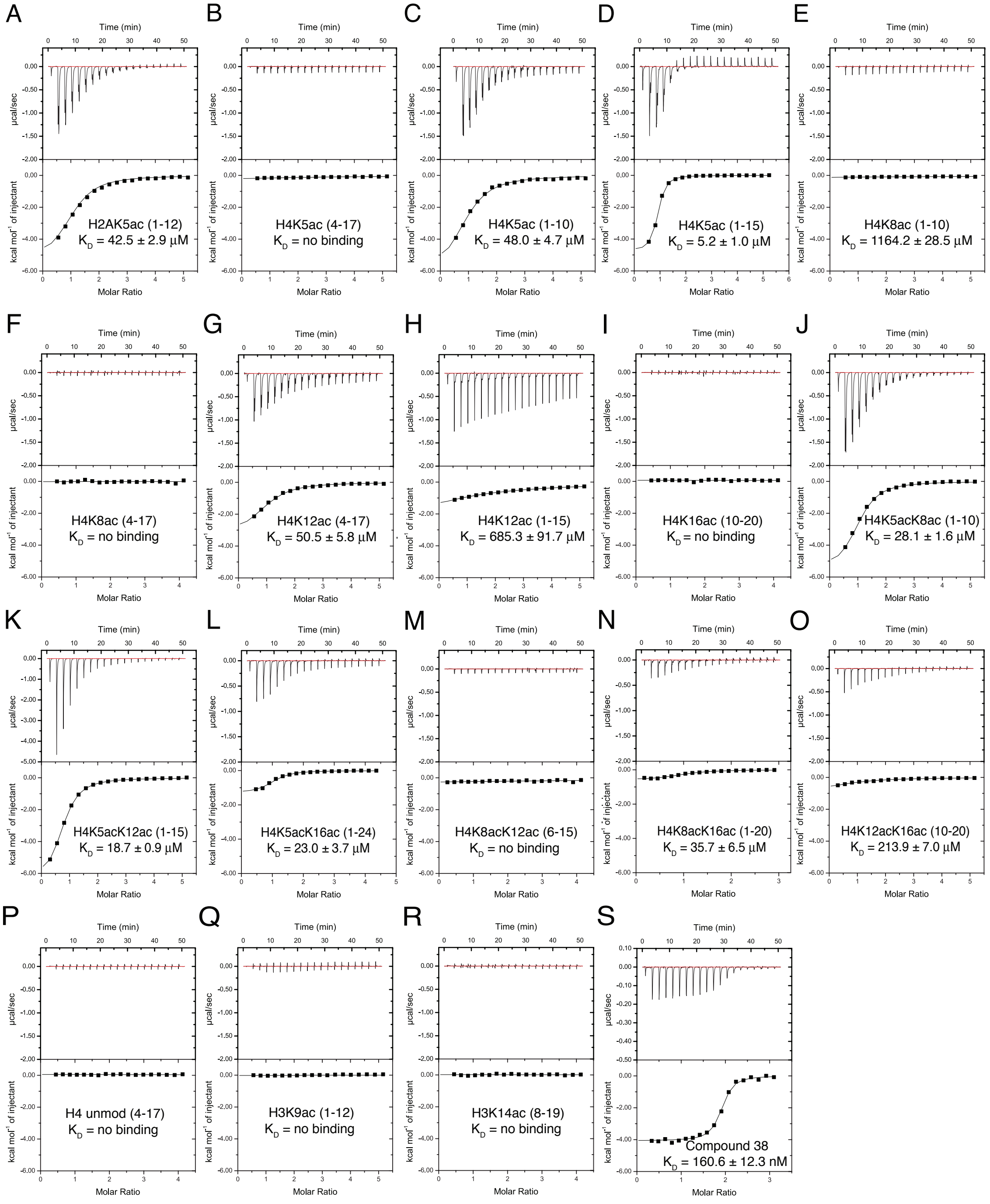

**Supplementary Figure 3. Interaction of the ATAD2B bromodomain with acetylated histone ligands.** (A) Superimposed $^{1}H$-$^{15}N$ HSQC spectra of the ATAD2B bromodomain, collected during titrating in the indicated histone peptides. The spectra are color-coded according to the protein/peptide ratio. (B) 2D $^{1}H$-$^{15}N$ HSQC spectra of $^{15}N$-labelled ATAD2B bromodomain with the complete HSQC assignments labeled.

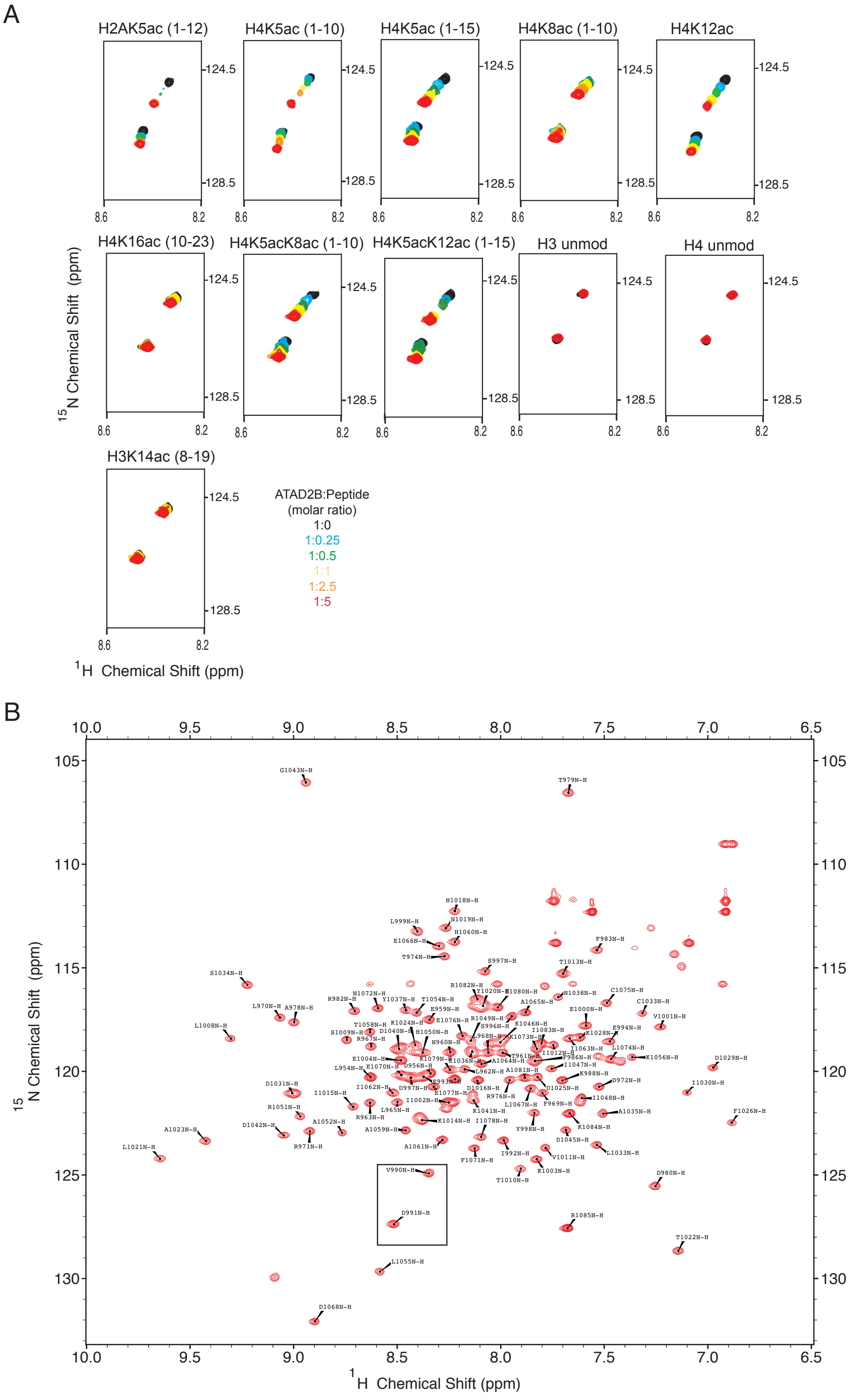

**Supplementary Figure 4.** *ATAD2* and *ATAD2B* gene structures and splicing patterns. (A) Sashimi plots of splice junctions between *ATAD2* and *ATAD2B* are shown. The number of reads spanning each junction is indicated by the size of the sashimi plot curve, and the number of reads spanning novel junctions is indicated by red sashimi plot curves. Coverage plots for total mapped reads are shown (total exon coverages are shown on the y-axis). (B) RT-PCR Analysis of ATAD2 and ATAD2B exon 21-22 splicing patterns.

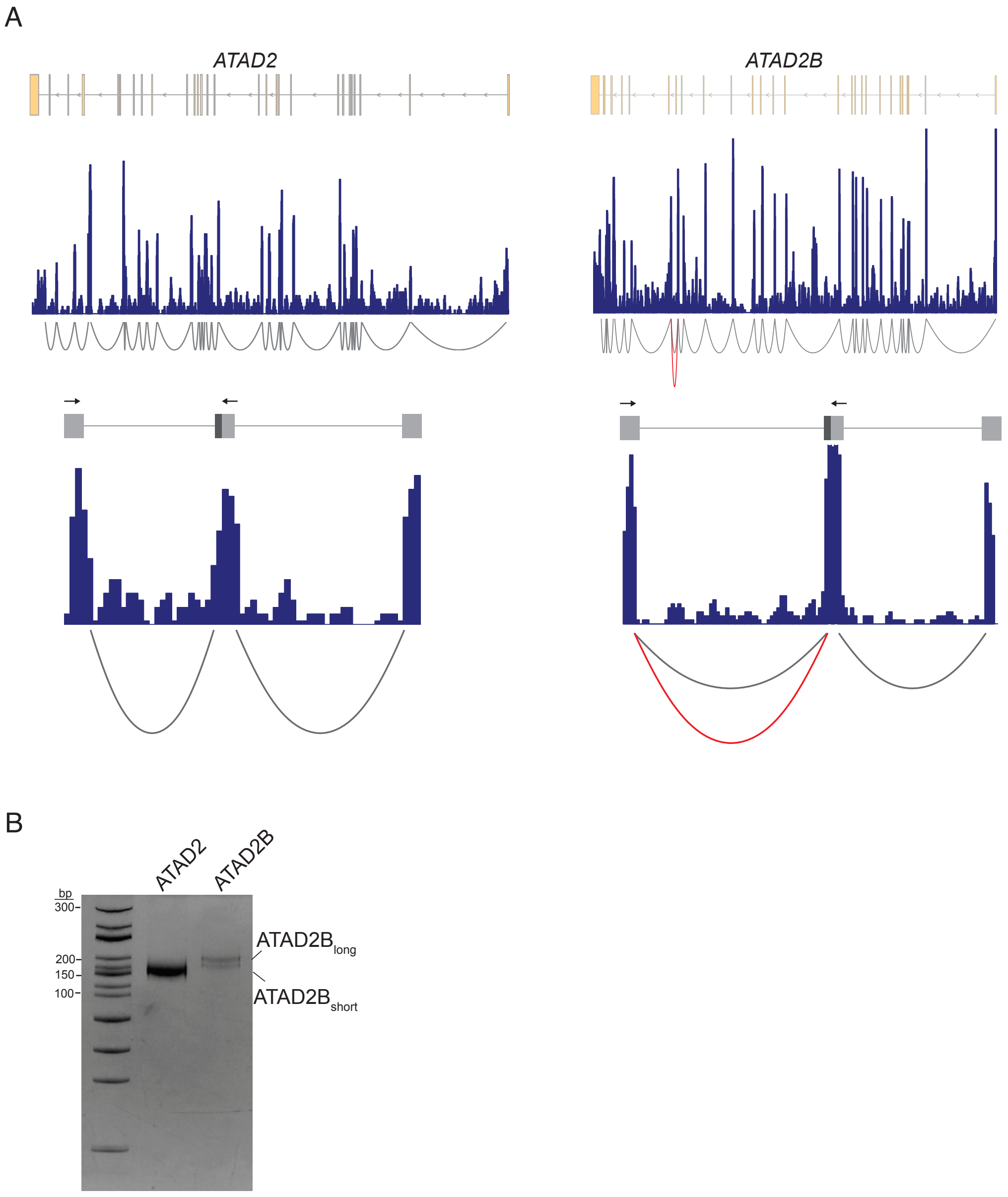

**Supplementary Figure 5. ITC measurements of the interaction between acetylated histone ligands and mutant ATAD2B bromodomain proteins.** (A-O) Exothermic ITC enthalpy plots for the binding of mutant ATAD2B bromodomain proteins to histone peptide ligands that are unmodified, mono- or di-acetylated. The calculated binding constants are indicated.

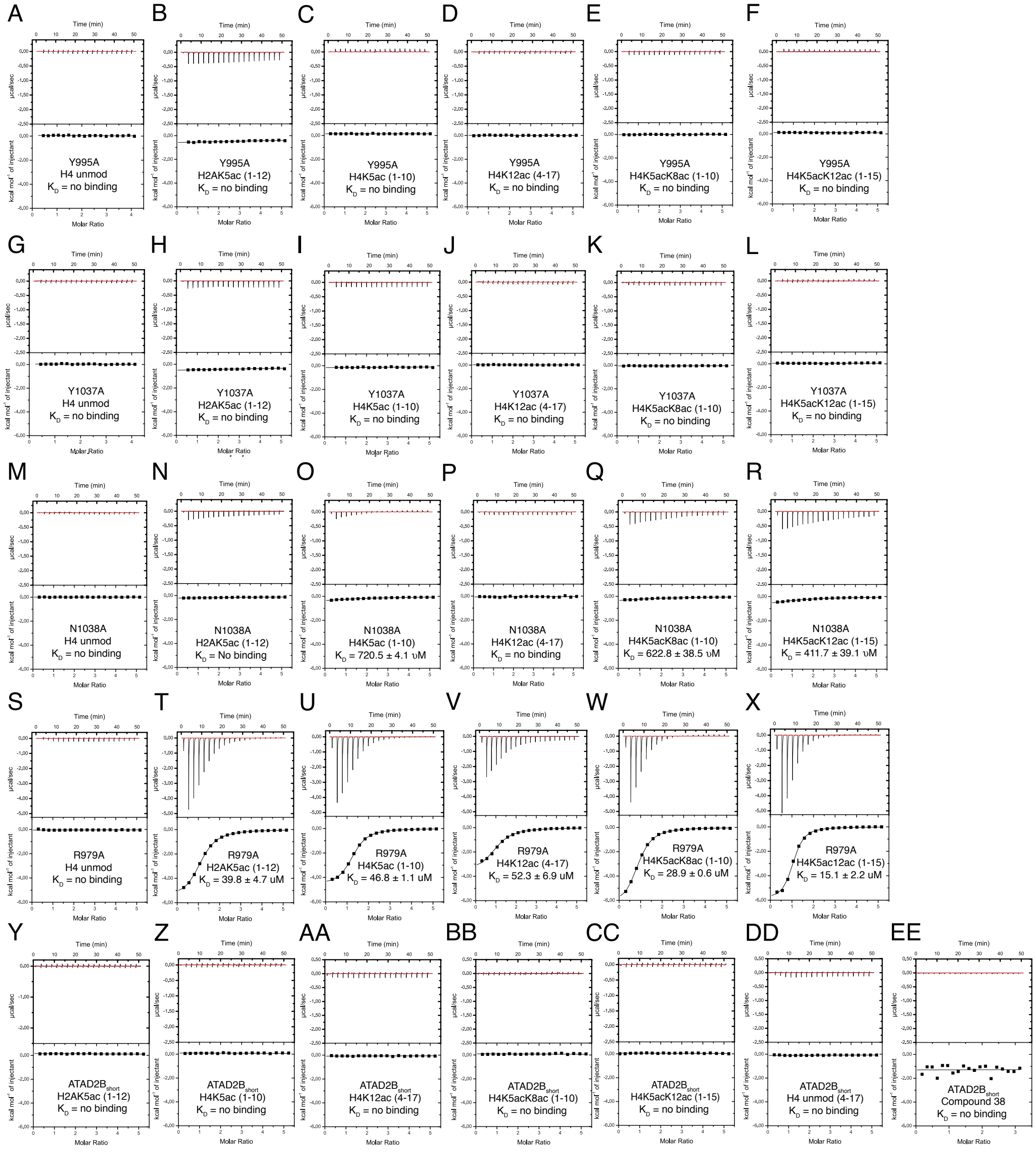

**Supplementary Figure 6.** Analysis of ATAD2B splice isoform expression from RNAseq data. (A) Scatterplot comparing the normalized counts of reads that map to ATAD2B (x-axis) and ATAD2B$_{short}$ (y-axis) isoforms across human breast cancer datasets from CCLE. Each dataset is colored by the breast cancer molecular subtype, the dashed line represents a 50:50 ratio of each isoform and the names of 4 cell lines with the highest expression of ATAD2B$_{short}$ are labeled. (B) A plot showing the ranked expression of ATAD2B: ATAD2Bshort across human breast cancer cell lines. (C) A box plot of the ATAD2B: ATD2B$_{short}$ ratios across human breast cancer molecular subtypes.

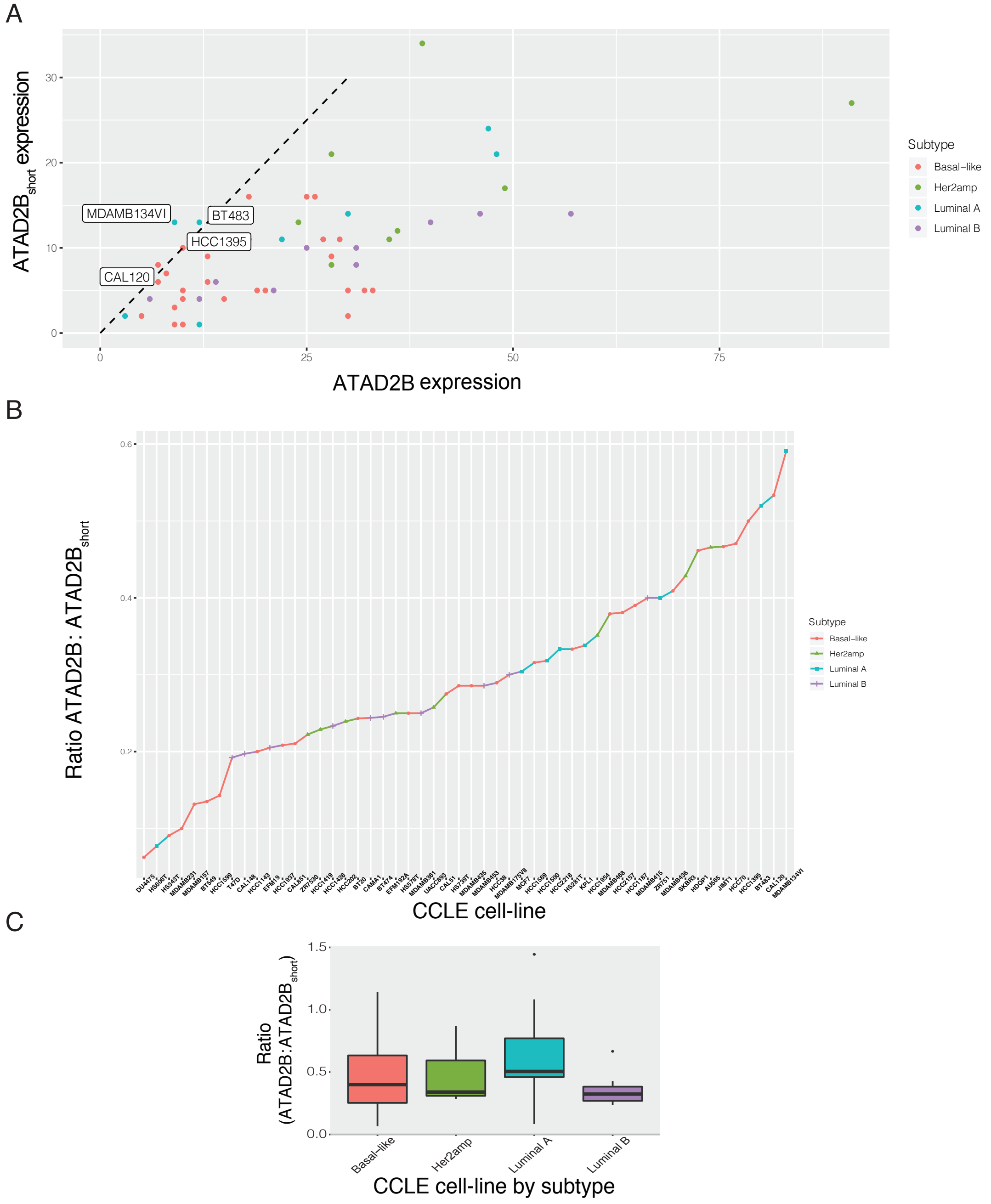

**Supplementary Figure 7. Sequence alignment of the ATAD2 and ATAD2B bromodomains.** Sequence alignment of the ATAD2 and ATAD2B bromodomain proteins corresponding to the amino acids in our ATAD2B bromodomain structure (residues 953-1085). Residues colored magenta are involved in hydrogen bonding to compound 38 for the ATAD2B bromodomain, and compound 42 for the ATAD2 bromodomain. Residues colored green are hydrophobic contacts. Residues that were mutated in this study are marked with a red vertical line. Sequence alignment was performed using the T-Coffee software [1].

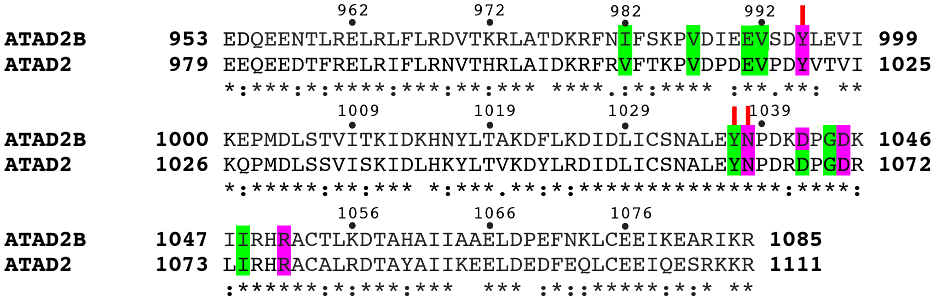



1. Notredame, C., Higgins, D.G. & Heringa, J. T-Coffee: A novel method for fast and accurate multiple sequence alignment. *J Mol Biol* **302**, 205-17 (2000).
